# Transport Dynamics of MtrD: an RND Multidrug Efflux Pump from *Neisseria gonorrhoeae*

**DOI:** 10.1101/2021.02.24.432727

**Authors:** Lauren Ammerman, Sarah B. Mertz, Chanyang Park, John G. Wise

**Affiliations:** Department of Biological Sciences, Southern Methodist University, Dallas, TX 75275; Center for Scientific Computation, Southern Methodist University, Dallas, TX 75275; Center for Drug Discovery, Design and Delivery; Southern Methodist University, Dallas, TX 75275; University of Texas Southwestern Medical Center, Dallas, TX, 75390

## Abstract

The MtrCDE system confers multidrug resistance to *Neisseria gonorrhoeae*, the causative agent of gonorrhea. Using free and directed Molecular Dynamics (MD) simulations, we analyzed the interactions between MtrD and Azithromycin, a transport substrate of MtrD and a last-resort clinical treatment for multidrug resistant gonorrhea. We then simulated the interactions between MtrD and Streptomycin, an apparent non-substrate of MtrD. Using known conformations of MtrD homologues, we simulated a potential dynamic transport cycle of MtrD using Targeted MD techniques (TMD), and we note that forces were not applied to ligands of interest. In these TMD simulations, we observed the transport of Azithromycin and the rejection of Streptomycin. In an unbiased, long-timescale simulation of AZY-bound MtrD, we observed the spontaneous diffusion of Azithromycin through the periplasmic cleft. Our simulations show how the peristaltic motions of the periplasmic cleft facilitate the transport of substrates by MtrD. Our data also suggest that multiple transport pathways for macrolides may exist within the periplasmic cleft of MtrD.

## Introduction

The gram-negative diplococcus *Neisseria gonorrhoeae* is responsible for the sexually transmitted infection (STI) Gonorrhea, and with over 87 million cases of Gonorrhea reported worldwide, antibiotic resistance in *N. gonorrhoeae* is a global health concern^*1, 2*^. Of these reported cases, approximately half are classified as drug resistant^*3*^. For multidrug resistant cases, only one recommended treatment remains – combination therapy with the antibiotics, ceftriaxone and azithromycin^*2*^. Gram negative pathogens like *N. gonorrhoeae* have evolved intricate mechanisms to overcome antimicrobial treatments, and among the most effective are the <u>r</u>esistance, <u>n</u>odulation, and cell-<u>d</u>ivision (RND) systems. The MtrCDE system is the sole RND system expressed *N. gonorrhoeae*, and modulates resistance to a variety of antibiotics^*4*^. The tripartite MtrCDE complex consists of the MtrD pump (uniprot Q51073), the MtrC adaptor (uniprot P43505), and the MtrE channel (uniprot Q51006)^*5–7*^. The overexpression and efflux activity of MtrCDE contribute significantly to clinical levels of antibiotic resistance in *N. gonorrhoeae*^*8, 7, 9*^. Importantly, both azithromycin and ceftriaxone are thought to be transport substrates of the MtrD pump.

MtrD is a critical agent of antibiotic resistance in *N. gonorrhoeae* and effluxes a variety of hydrophobic and amphiphilic substrates^*6, 8*^. The pump assembles as a homotrimer, with each monomer consisting of a periplasmic domain and 12 transmembrane helices^*5, 10*^ (Figure 1A,B). The periplasmic domain contains the “periplasmic cleft” (or “cleft”) that facilitates substrate transport, and a “docking” domain that interfaces with MtrCE (Figure 1B). The periplasmic cleft is divided into the Access Pocket (PC1 and PC2 domains) and the Deep Pocket (PN1 and PN2), are alternately the proximal and distal multidrug binding sites (Figure 1C). Access to the Deep Pocket and distal site is controlled by the flexible and highly conserved Gate-Loop, or “G-Loop” (Figure 1C, magenta). The transmembrane domain houses the highly conserved Proton Relay Network (PRN)^*5*^ (Figure 1D). Changes in the protonation states of the PRN correlate with vertical shearing of the TM helices and the peristaltic motions of the periplasmic cleft^*11, 12*^.

**Figure 1.**
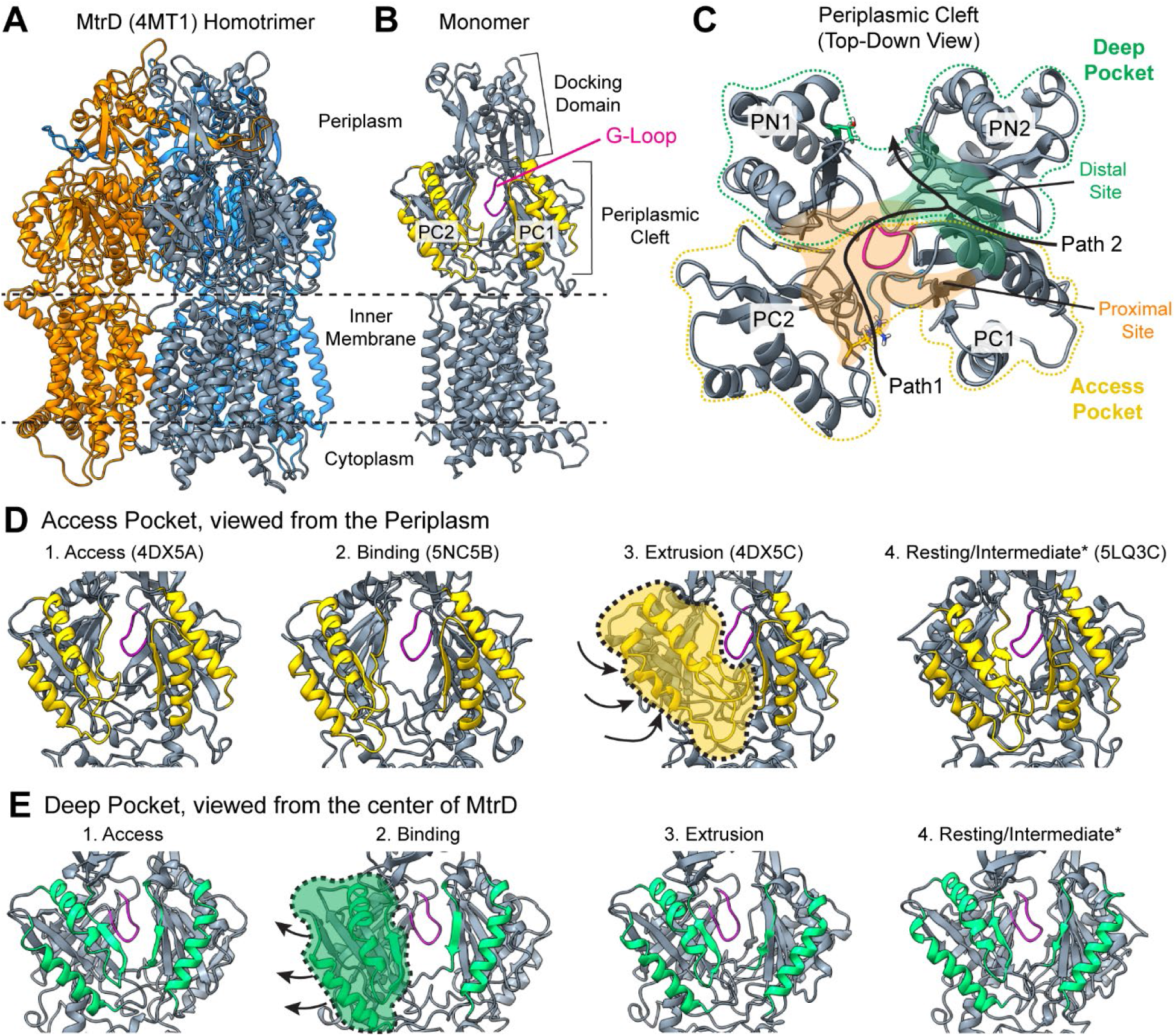
The MtrD Efflux Pump from *Neisseria gonorrhoeae*. **(A)** The MtrD homotrimer with subunits colored orange, gray and blue. **(B)** An MtrD monomer with helices of the Access Pocket in yellow and the G-Loop in magenta. **(C)** The periplasmic cleft viewed as if looking from the periplasm towards the inner membrane; helices of the Access Pocket in yellow, helices of the Deep Pocket in green, G-Loop in magenta. K823 and R714 may contribute to macrolide recognition (in orange sticks), and F612 and F610 may facilitate substrate selectivity^*10*^. **(D)** The Access Pocket viewed from the periplasm; arrows show how PC2 (shaded yellow) closes during the transport cycle. **(E)** The Deep Pocket viewed from the central pore; arrows show how PN2 (shaded green) opens during the transport cycle. Stages of the transport cycle are labeled Access (also ‘Loose’), Binding (‘Tight’), Extrusion (‘Open’), and Resting/Intermediate with the corresponding crystal structure of the MtrD homologue AcrB (5NC5, 4DX5) or CmeB (5LQ3) in parentheses^*5, 15, 16, 19*^. Helices of the Access or Deep Pockets are colored differentially to aid in the visualization of conformational changes during the transport process.

Three conformations of the transport cycle have been identified: (1) “Access/Loose”, (2) “Binding/Tight” (3) “Extrusion/Open”^*10, 13–16*^ (Figure 1E-F, S1). A fourth putative structure, “Resting”, has also been identified (Figure 1E, F). Substrate-free MtrD adopts a symmetrical conformation with each monomer in Access^*5*^. Upon the binding of a transport substrate, each monomer cycles from Access, to Binding to Extrusion - with the ‘Resting’ state as a potential last step - in a functional rotation mechanism^*11*^. During this process, the substrate is somehow extruded through the periplasmic cleft^*10, 13*^. Substrate movement is thought to occur in three stages: substrates 1) bind to the proximal site in the Access Pocket, (2) bind to the distal site in the Deep Pocket, and 3) are released into the funnel formed by the docking domains of the homotrimer^*5, 10 17*^. Small hydrophobic substrates may bypass step 1 and enter the Deep Pocket directly through the cleft formed by PC1 and PN2. Thus, the major conformational states of RND pumps and potential binding locations of substrates have been identified largely through X-ray crystallography and cryo-EM^*10, 12, 13, 15, 17*^. However, it is unclear how these conformational changes facilitate the movement of transport substrates through the periplasmic cleft.

To understand the process of substrate transport, we simulated a putative efflux cycle of MtrD using the known conformations of MtrD homologues and Targeted Molecular Dynamics (MD) techniques. In our Targeted MD simulations, a putative substrate or non-substrate of MtrD was included. Forces were applied to α-carbons of the MtrD backbone, but not to transport ligand(s) of interest, or to any other atoms in the system. MtrD substrates are thought to first bind within the proximal site to initiate transport^*5*^, but as shown in Figure 1C, the proximal site is large and contains residues from both the Access and Deep Pockets. Consequently, we chose two locations within the proximal site for Targeted MD simulations of ligand transport. To complement our biased simulations, we also performed an unbiased 1.5 μs MD simulation of substrate bound MtrD in the Access state. Lastly, we analyzed the molecular landscape of the periplasmic cleft.

In our Targeted MD simulations, we observed the transport of the macrolide antibiotic azithromycin (AZY). AZY was observed to take an alternate transport pathway that bypassed the distal site, and while these results were unexpected, we found that clinical data supports the existence of this alternate route^*18*^. Including a fourth “Intermediate/Resting” structure^*16*^ increases the transport distance of AZY, suggesting that this state might represent a “re-setting” of the monomer at the end of the efflux cycle. We also observed the rejection – or retention – of streptomycin (SRY) from the proximal / Access site, supporting the hypothesis that SRY is not a transport substrate^*6*^, nor a likely inhibitor, of MtrD. In our unbiased simulation of AZY-bound MtrD in Access, we observed the spontaneous movement of AZY from the Access Pocket to the Deep Pocket. These data support the hypothesis that substrate diffusion into the Deep Pocket occurs slowly in the absence of changes in the PRN^*11*^. Lastly, we found that access to areas with specific molecular signatures is alternately allowed and restricted by conformational changes of the periplasmic cleft. Taken together, our data suggest that multiple transport pathways through the periplasmic cleft may exist – even for large macrolides like AZY.

## Results

### Modeling Substrate Transport with Targeted MD Simulations

Using Targeted MD techniques as described in^*20*^, we applied small forces to the α-carbons of MtrD to model the conformational changes of a putative transport cycle (Figure 1D-E, S1). Structures of MtrD homologues in Access, Binding, Extrusion, and Resting/Intermediate were used as targets (See Methods, Figure S1). As ligands, we chose azithromycin (AZY) and streptomycin (SRY)^*6, 8*^. AZY is one of the last remaining therapeutics for multidrug resistant gonorrhea and a transport substrate of MtrD^*8*^. Mutations in MtrD have been shown to modulate AZY resistance in *N. gonorrhoeae*^*18, 21, 22*^. Thus, it is of interest to understand how MtrD transports substrates of this size and clinical importance. SRY is one of the few recognized non-substrates of MtrD and is of a comparable size to AZY (SRY: ~581 Da, AZY: ~749 Da). Thus, SRY is an appropriate choice as a negative control for transport in our MD simulations.

Large substrates are thought to first bind at the proximal site in the Access Pocket, and then bind at the distal site in the Deep Pocket (Figure 1C, Path 1)^*10, 17*^. But both SRY and AZY are too big to use the alternate entrance for small substrates (Figure 1C, Path 2). The precise binding location of AZY or SRY in the proximal binding site is unclear^*10*^. To generate reasonable ligand starting positions for MD simulations, we independently docked AZY and SRY to the periplasmic cleft of MtrD using *Autodock Vina*^*5, 23, 24*^ (Figure S2). Predicted docking poses for both SRY and AZY clustered near the G-Loop, with outliers at the entrance or exit of the cleft (Figure S2). These docking calculations show that MtrD in “Access” can accommodate multiple binding modes of AZY and SRY despite their large sizes.

From these initial experiments, we selected two positions within the proximal site to use as “start sites” for MD simulations. At Site 1, SRY and AZY are interacting with the foremost residues of the proximal site (Figure 2A, Site “1”). Here we tested whether SRY, a putative non-substrate of MtrD, will be retained within the Access Pocket. The foremost residues of the Access Pocket are thought to serve as a selectivity filter; thus, we would expect AZY to be retained at this site. However, Targeted MD simulations use relatively short timescales, and the process of substrate association with the G-Loop is thought to occur slowly with the transporter in the Access conformation^*15*^. Thus, in our simulations from Site 2, which is located deeper within the access pocket, SRY and AZY are associating with the proximal site and the G-Loop (Figure 2A, “2”). Here we test if the conformational changes of MtrD result in the transport of a known substrate (AZY) or a suspected non-substrate (SRY). We performed 20 independent Targeted MD simulations per ligand at Sites 1 and 2 (n = 20 Targeted MD simulations per ligand, per protonation state, per site). Forces were not applied to AZY or SRY.

**Figure 2.**
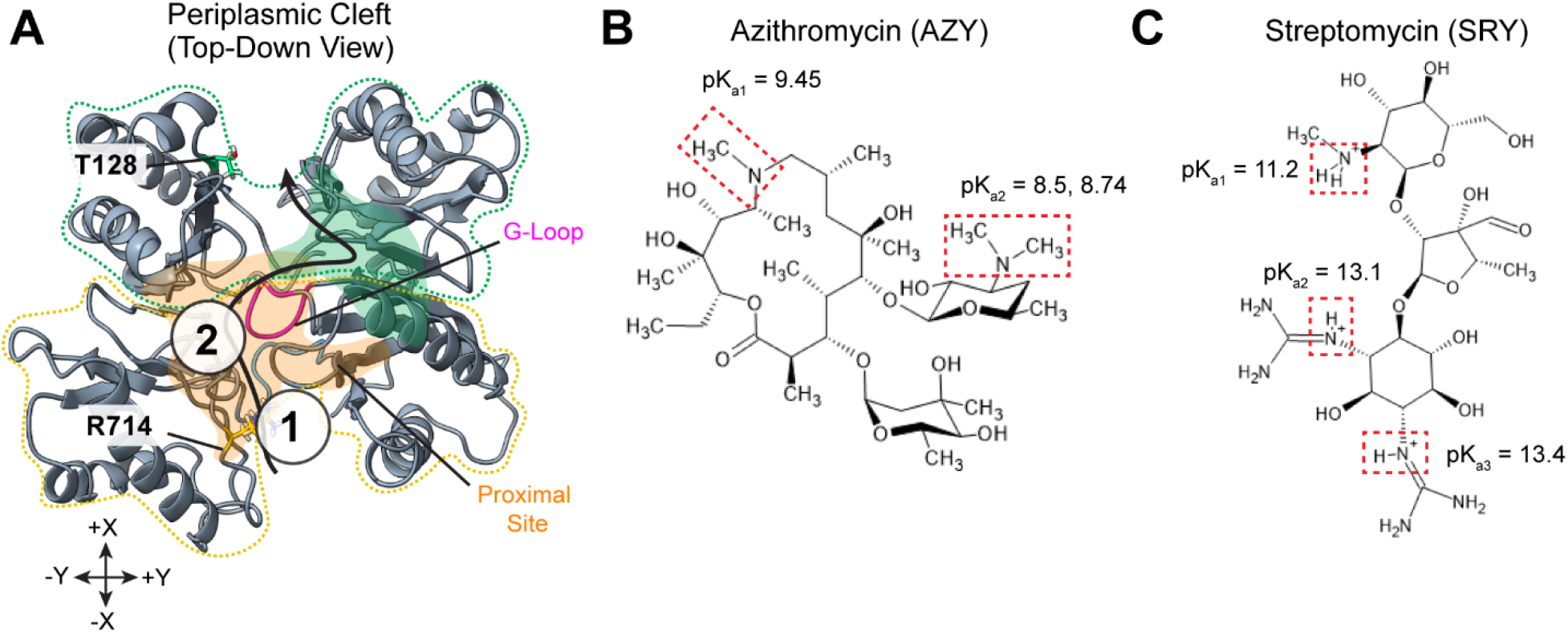
The Ligand Starting Sites within the Periplasmic Cleft of MtrD. **(A)** The periplasmic cleft of MtrD. The Deep Pocket (green) and Access Pocket (yellow) are outlined in dashed lines. The proximal and distal binding sites are shaded orange and green, respectively. The putative pathway for large substrates is shown as a black arrow, moving from the proximal site, towards the distal site. The G-Loop is shown in Magenta. R714 and T128 – residue “checkpoints” for measuring substrate movement – are labeled. Start sites 1 and 2 are shown as numbered circles. Estimated binding affinities are as follows; −7.7 kcal/mol for AZY at Site 1; −7.4 kcal/mol for SRY at Site 1; −7.4 kcal/mol for AZY at Site 2; −7.3 kcal/mol for SRY at Site 2. **(B)** The structure and pK_a_ values of ionizable amine groups in Azithromycin (AZY)^*21, 25*^. **(C)** The structure and pK_a_ values of ionizable groups in Streptomycin (SRY)^*26, 27*^.

### Each Protonation State of AZY is Included in Targeted MD Simulations

MtrD transports diverse hydrophobic and amphiphilic substrates^*7, 8, 10*^. At physiological pH ranges (between 7 and 7.5), three potential protonation states of the MtrD substrate AZY are possible: uncharged AZY_neu_ (~1% probability); singly protonated, positively charged AZY_h1_ (~5-8% probability); and doubly protonated, positively charged AZY_h2_ (~92-95% probability) (Figure 2B). The aforementioned probabilities were calculated based upon experimentally derived pKa values for each protonation site^*21, 28*^ of AZY (pK_a_1 9.45, is and pK_a_2 is 8.5-8.74); however, we note that pK_a_ values are sensitive to changes in the surrounding environment. *N. gonorrhoeae* colonizes areas of the human body with a variety of pH ranges^*2, 29, 30*^. Furthermore, it is thought that the neutral, unprotonated form of AZY might diffuse most readily through the bacterial membrane, and thus encounter MtrD in this state^*25*^. Finally, transport ligands are also exposed to the microenvironment of the periplasmic cleft. Thus, MtrD might encounter each protonation species of AZY. To investigate how each species of AZY might interact with MtrD, we tested each protonation state in Targeted MD simulations (see Methods). We note that only one protonation state of SRY (3+) is likely to occur at physiological pH ranges (Figure 2C).

### AZY is Retained in the Access Pocket during Simulations from Site 1

Due to its size and the timescale of our Targeted MD simulations (see Methods), AZY was not expected to reach the G-Loop when started at Site 1 (Figure 3A). However, if the foremost residues of Access Pocket act as a filter, then we would expect that AZY – a known substrate of MtrD^*9, 18, 21, 22*^ – would be retained at this site. Consistent with these expectations, AZY remained between the PC1 and PC2 domains of the Access Pocket during Targeted MD simulations from Site 1 (Figure 3B). In 2/20 simulations, AZYneu slipped into the solvent but remained associated with exterior of the periplasmic cleft. This occurred in 1/20 simulations for AZY_h1_, and 1/20 for AZY_h2_. This behavior is unsurprising given the fact that the periplasmic cleft closed before AZY could reach the G-Loop. These data suggest that diffusion through the Access Pocket occurs slowly. Full simulation results are shown in Figure S5 and Movie S1 [5].

**Figure 3.**
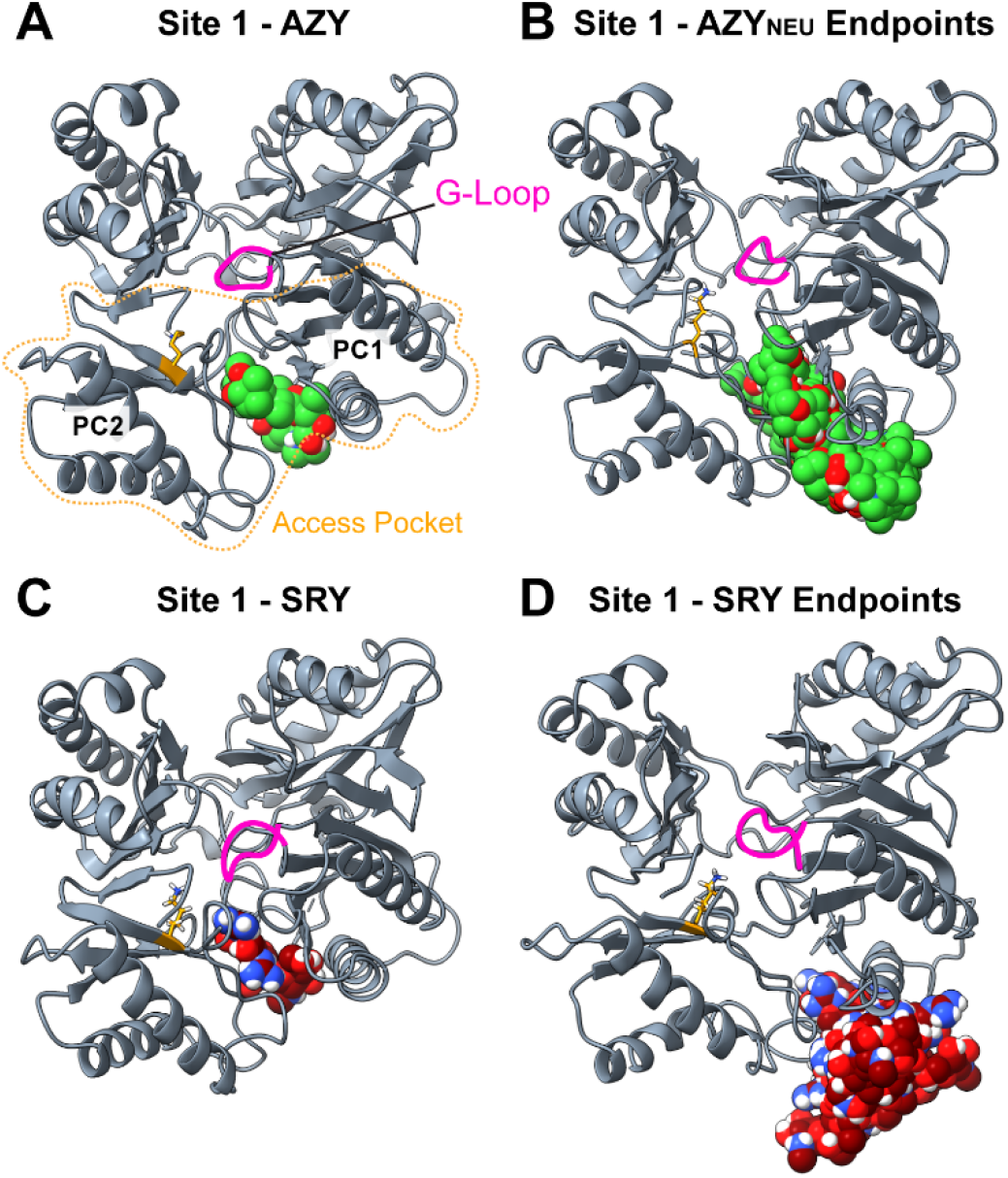
Targeted MD Simulations of SRY and AZY from Site 1. **(A)** shows AZY_NEU_ (green) in its starting position for simulations from Site 2. The Access Pocket is outlined in yellow and its domains are labeled. The G-Loop is shown in Magenta. **(B)** shows all ending poses of AZY_NEU_ from Site 2. **(C)** shows SRY (maroon) in its starting position for simulations from Site 2. **(D)** shows all ending poses of SRY from Site 2 simulations. Note that SRY has exited the Access Pocket in 20/20 simulations.

### SRY is Rejected from the Access Pocket during Simulations from Site 1

SRY is not a recognized substrate of MtrD; thus, we examined our simulations from Site 2 to see whether SRY could successfully enter the periplasmic cleft (Figure 3C)^*6*^. In contrast to the observed behavior of AZY, SRY was observed to move ‘backwards’, i.e. away from the interior of the periplasmic cleft, and into the solvent (Figure 3D). Once SRY exited the periplasmic cleft, it remained closely associated with D709 on the cleft exterior in 14/20 trajectories (>70% of the simulation time). We note that SRY and AZY docked to this location with comparable estimated affinities: −7.7 kcal/mol for AZY and −7.4 kcal/mol for SRY. Thus, the difference between the behavior of SRY and AZY cannot be attributed to a large discrepancy between estimated binding affinities. These data support the hypothesis that residues of the Access Pocket act as a selectivity filter, and support the hypothesis that SRY is not a transport substrate of MtrD^*6*^. Representative trajectories of SRY at Site 1 are shown in Movie S2.

### AZY is Transported by MtrD in Targeted MD Simulations from Site 2

Our Targeted MD simulations of AZY from Site 1 confirmed that AZY is indeed retained at the Access Pocket, but these simulations did not provide enough time for AZY to associate with the G-Loop. Therefore, we further examined the simulation results of AZY from Site 2 for transport behavior. These simulations from Site 2 modeled a putative stage of transport in which AZY has diffused through the Access Pocket and associated fully with the G-Loop.

#### Defining “Transport” in Targeted MD Simulations

It was of interest to quantify substrate movement – if any – during our simulations. Ligand movement was first quantified by calculating the root mean squared deviation (RMSD) of the ligand from its starting position over time. However, since RMSD does not necessarily distinguish translational movement from rotational movement, we also calculated the distance from the ligand’s center of mass (LigandCOM) to “checkpoint” residues within the interior of the periplasmic cleft. We chose the α-carbon of R714 at the entrance of the periplasmic cleft, and the α-carbon of T128 at the exit of the cleft (Figure 2A). If a ligand is transported in TMD simulations, the LigandCOM to R714_α-carbon_ distance should increase, and the Ligand_COM_ to T128_α-carbon_ distance should decrease (Figure 2A). Thus, in our simulations, we defined a ligand as “transported” if, by the end-of-simulation, each of the following conditions were met: the Ligand_COM_ to R714_α-carbon_ distance was greater than or equal to 18 Å, the Ligand_COM_ to T128_α-carbon_ distance was less than or equal to 15 Å, and the RMSD_ligand_ was at least 8.5 Å.

#### Transport of AZY by MtrD in Targeted MD Simulations

Based upon the previously defined distance cutoffs between LigandCOM (center of mass) and R714_α-carbon_ or T128_α-carbon_, simulation outcomes were divided into two clusters: “Transported” (Figure 4A-C) versus “Non-Transported” (Figure S4). In the “Transport” cluster of trajectories, AZY was observed to move from the Access Pocket, past the G-Loop, and into the Deep Pocket. AZY then continued to travel along the PN1 domain in the Deep Pocket until it reached the exit of the periplasmic cleft and was positioned for dissociation (or release) into the funnel domain. The following frequencies of transport were observed: AZY_neu_ was transported in 12/20 simulations; AZY_h1_ was transported in 3/20 simulations; AZY_h2_ was transported in 6/20 simulations. Since MtrD is a member of the hydrophobe-amphiphile efflux family of RND transporters, it is unsurprising that the most lipophilic species – AZY_neu_ – exhibited the highest frequency of transport in TMD simulations. In the “Non-Transport” cluster of AZY trajectories, two possible outcomes were observed: 1) AZY remained straddling the G-Loop, or 2) AZY moved into the Deep Pocket, but did not travel fully towards the exit of the cleft (Figure S4). Examples of a both Transport and Non-Transport trajectories are shown in Movie S3.

**Figure 4.**
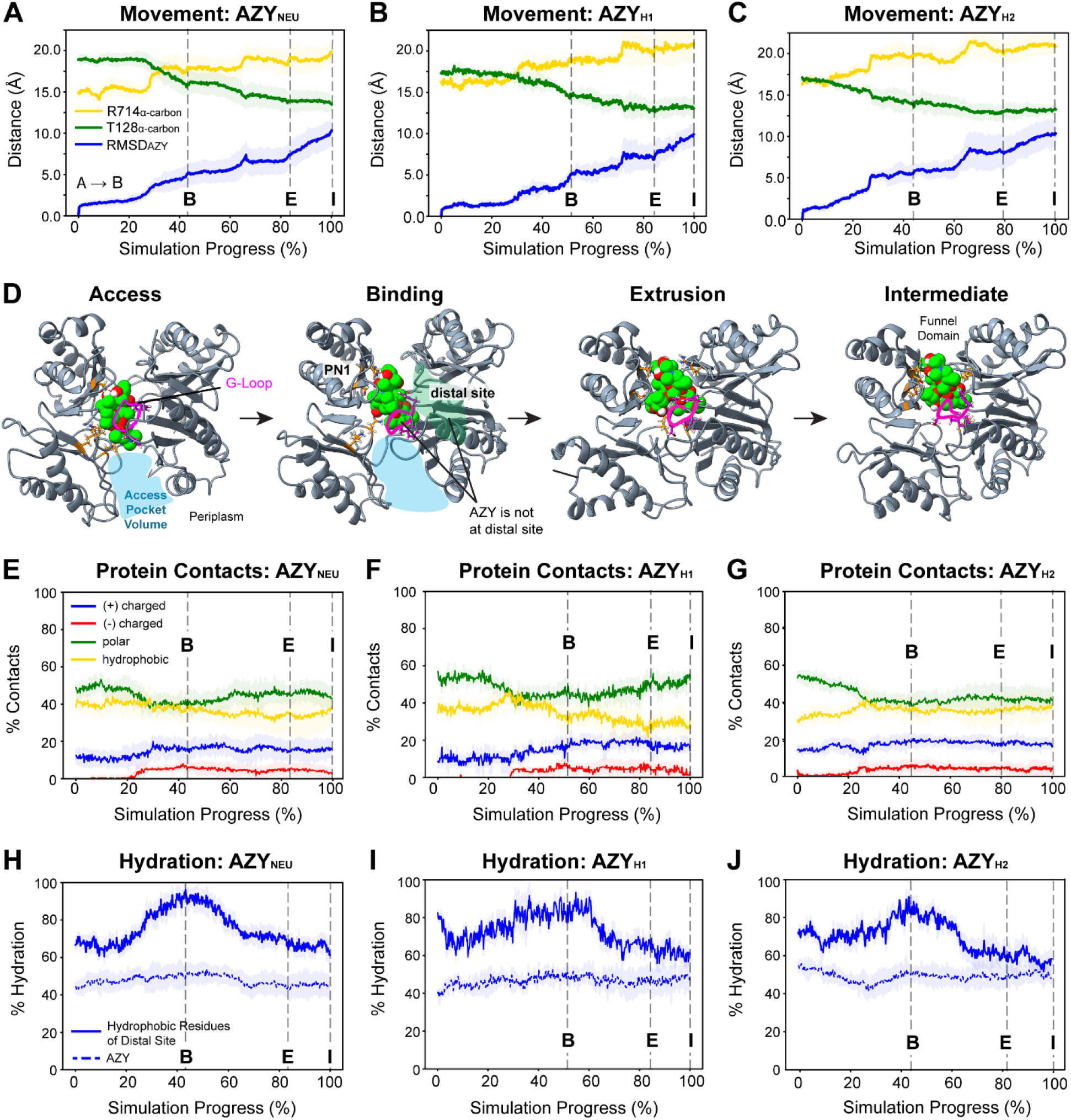
Transport of AZY by MtrD in Targeted MD Simulations from Site 2. **Panels (A-C)** show the RMSD_AZY_, AZY_COM_-T128_α-carbon_ distance, or AZY-R714_α-carbon_ distance in Angstroms (Å) over time. **(D)** shows transport of AZY_neu_ (lime) in a representative trajectory. A blue overlay shows the approximate size of the Access Pocket when open to the periplasm. The G-Loop is magenta, and the distal site is shaded green. **(E-G)** show the chemical nature of MtrD-AZY contacts over time; the majority are either polar or hydrophobic. **(H-J)** show the hydration of the SASA_AZY_, or of the SASA_distal hydrophobic_, over time. Data in **(A-C),** and **(E-J)** show the mean ± one standard deviation in shading. Dashed grey lines mark when MtrD reaches a structural checkpoint, starting with MtrD in Access (“A”), and moving from Binding (“B”), to Extrusion (“E”), to Intermediate/Resting (“I”). Timepoints between the dashed lines indicate that MtrD is transitioning between two states.

### AZY did not Bind to the Distal Site in Targeted MD Simulations

The distal site is composed of primarily hydrophobic residues and is formed by the PN2 and PC1 domains (Figure S3). The cryo-EM structure of MtrD_CR103_, an MtrD variant which confers elevated AZY resistance, shows the macrolide erythromycin (ERY) bound at the distal site when MtrDCR103 is in the Binding conformation (6VKT)^*10*^. In this orientation, we calculated that 43% of MtrDCR103-ERY contacts were contributed by PN2, 43% by PC1, and 14% by the G-Loop (calculated with ChimeraX). Based upon these data and structures of ERY with AcrB, the distal site is thought to be the “second stop” on a ligand’s transport pathway (with the proximal site as the “first stop”), and we expected AZY to interact with the distal site when MtrD adopted the Binding conformation^*8, 10, 17, 31*^. During the subsequent transition from Binding to Extrusion, we then expected AZY to move along PN2 towards the exit of the cleft.

Unexpectedly, we found that AZY did not bind at the distal site when MtrD approached or adopted the Binding conformation (Figure 4D, “Binding”). If AZY was transported, it instead moved along an alternate route mediated by PN1. From Binding onward ≥ 40% of MtrD-AZY contacts were contributed by PN1. In contrast, PN2/PC1 – which form the distal site – accounted for a smaller share (6 to 15% and 0.6 to 7%, respectively) (Supplemental Table 2). Hydrophobic interactions at Binding can be attributed to the G-Loop, and many of the polar interactions to PN1 (Figure 4E-G, Table 1). If AZY was not transported, it still did not bind at the distal site, and PN1 still contributed the largest share of contacts (Figure S5, Table S2). Our data suggest that the distal site may not be an obligatory or essential “second stop” during transport.

**Table 1.** MtrD-AZY Contacts by Domain of the Periplasmic Cleft in a Long Timescale Simulation. Contacts are residues whose α-carbon is within 4 Å of AZY at each timepoint of the 1simulation. Individual domains of the periplasmic cleft are as defined in^*10*^ or as indicated. Contact analysis was performed with scripts in *Tcl* and python.

|  |  | Percent (%) of Protein-Ligand Contacts Contributed by Domain or Region |  |  |  |  |  |  |  |
| --- | --- | --- | --- | --- | --- | --- | --- | --- | --- |
|  | Simulation Checkpoint | G-Loop | PC1 | PC2 | PN1 | PN2 | F-Loop | AP (PC1 + PC2) | DP (PN1 + PN2) |
| AZY<sub>neu</sub> | 0% | 12 | 12 | 34 | 36 | 1 | 4 | 50 | 37 |
|  | 20% | 16 | 5 | 35 | 44 | 0 | 0 | 50 | 44 |
|  | 40% | 18 | 4 | 35 | 38 | 5 | 0 | 39 | 43 |
|  | 60% | 15 | 7 | 34 | 37 | 7 | 0 | 41 | 44 |
|  | 80% | 14 | 18 | 14 | 40 | 12 | 3 | 35 | 52 |
|  | 100% | 11 | 17 | 14 | 44 | 12 | 3 | 34 | 55 |

### AZY Travels Farther when a Fourth “Intermediate” Structure is Included

A fourth potential step of the catalytic transport cycle was identified by Su et al. in CmeB from *Campylobacter jejuni*, which shares a 39% sequence identity with MtrD^*16*^. In this fourth “Resting” structure, the Access Pocket is closed to the periplasm, and the orientation of the TM helices mimics that of the Extrusion conformation (Figure 1E-F, “Resting/Intermediate”)^*16*^. Su et al. postulated that this structure could be a conformational intermediate and a low energy resting state for the pump, and found it was most stable in the absence of a proton gradient^*16*^. However, a proton gradient across the plasma membrane is almost always present in actively metabolizing gram negative bacteria^*32*^. If the longevity of the Resting structure depends upon the absence of a proton gradient, then it is unlikely to serve as a low energy resting state for the transporter. Nonetheless, this conformation was observed in a crystal structure of CmeB (5LQ3) and in sm-FRET experiments^*16*^.

Based upon the orientation of its TM helices and Access Pocket, we alternatively hypothesized that the Resting structure is a conformational intermediate between Extrusion and Access. Thus, the Resting conformation was included as the last stage in Targeted MD simulations. We found that including the Resting structure in the target sequence resulted in the additional movement of AZY in the transport direction (Figure 4A-C, “E → I”). This increased movement appears to result from the additional constriction of the Deep Pocket during the transition from Extrusion to Intermediate^*16*^. In addition to serving as a conformational “re-set” from Extrusion to Access, the Resting structure may also help position particularly bulky substrates like AZY for dissociation from MtrD. We therefore refer to this fourth structure as the “Intermediate” (Figure 4, abbreviated as “I”).

### Substrate Hydration of Mediates Transport of AZY from Site 2

It has been suggested that water mediates the diffusion of transport substrates through the periplasmic cleft^*31*^. To elucidate the role of water in substrate transport by MtrD, we calculated the percent hydration of the solvent accessible surface area of AZY (SASA_AZY_) over the course of the Targeted MD simulations (Figure 4H-J). If water was indeed mediating substrate transport through the cleft, we would expect a constant and significant level of substrate hydration throughout the simulations. To ascertain if close hydrophobic interactions between AZY and the distal site were occurring, we also calculated the percent hydration of the hydrophobic residues in the distal site (SASA_distal hydrophobic_) (Figure S3). If AZY was found to be closely interacting with hydrophobic residues of the distal site, then the percent hydration of the SASA_distal hydrophobic_ should decrease as MtrD approaches or adopts the Binding conformation.

When AZY was classified as “Transported”, the SASA_AZY_ was ≥ 45% hydrated throughout the simulations. When MtrD was in the Binding conformation, when AZY was expected to bind at the distal site, the SASA_distal hydrophobic_ exhibited a 20% increase (up to 80-90%) (Figure 4H-J). This increase in hydration of the distal site can be attributed to 1) the lack of AZY bound at this location, and 2) the wide-open state of the Access Pocket, which allows an influx of water into the cleft. Analyses of MtrD-AZY contacts by protein domain confirm that significant interactions between AZY and the distal site (PC1/PN2) did not occur as MtrD approached or adopted the Binding structure (Table S2). Notably, the SASA_distal hydrophobic_ was ≥ 60% hydrated regardless of simulation outcome (Figure 4H-J, Figure S4D-F). Our data support the hypothesis that water mediates substrate movement through the periplasmic cleft^*31*^.

### AZY Simulations at Site 2 Diverge in the Transition from Access to Binding

Using analyses of MtrD-AZY contacts throughout the simulations, we found that the “Transport” trajectories diverged from the “Non-Transport” trajectories during the transition from Access to Binding (Supplemental Table 2). If AZY successfully slipped past the G-Loop during this period, then AZY was squeezed towards the exit as the Access Pocket subsequently closed. Notably, this movement of AZY – past the G-Loop and into the Deep Pocket – occurred during the *transition* from Access to Binding, not when MtrD had fully adopted the Binding structure. Thus, AZY can diffuse past the G-Loop even if the cleft is not in its most open conformation.

### Streptomycin was not transported in Targeted MD Simulations from Site 2

In Targeted MD simulations with SRY started at Site 2, using the same methodology and transport criteria as was used with AZY, SRY was classified as ‘not transported’ in 20 out of 20 trajectories (Figure 5). The mean RMSD_SRY_ was 6.2 ± 1.7 Å (Figure 5A). The starting and ending positions of SRY from Site 2 are shown in Figure 5C and Movie S4. We observed that SRY remains closely associated with the G-Loop, particularly with F612 and S613, throughout the simulations. 40% of the SASA_SRY_ was consistently hydrated (Figure 5D). Polar contacts with PN1 and PC2 contributed the majority of MtrD-SRY interactions throughout the simulated conformational cycle, followed by hydrophobic contacts with F612 of the G-Loop (Figure 5C,E and Supplemental Table 2). SRY was not observed to interact significantly with PC1 or PN2 (Supplemental Table 2). We also observed that SRY displayed significantly more conformational flexibility than AZY. Our results indicate that SRY, if placed in the proximal binding site at Site 2, does not exhibit transport behavior.

**Figure 5.**
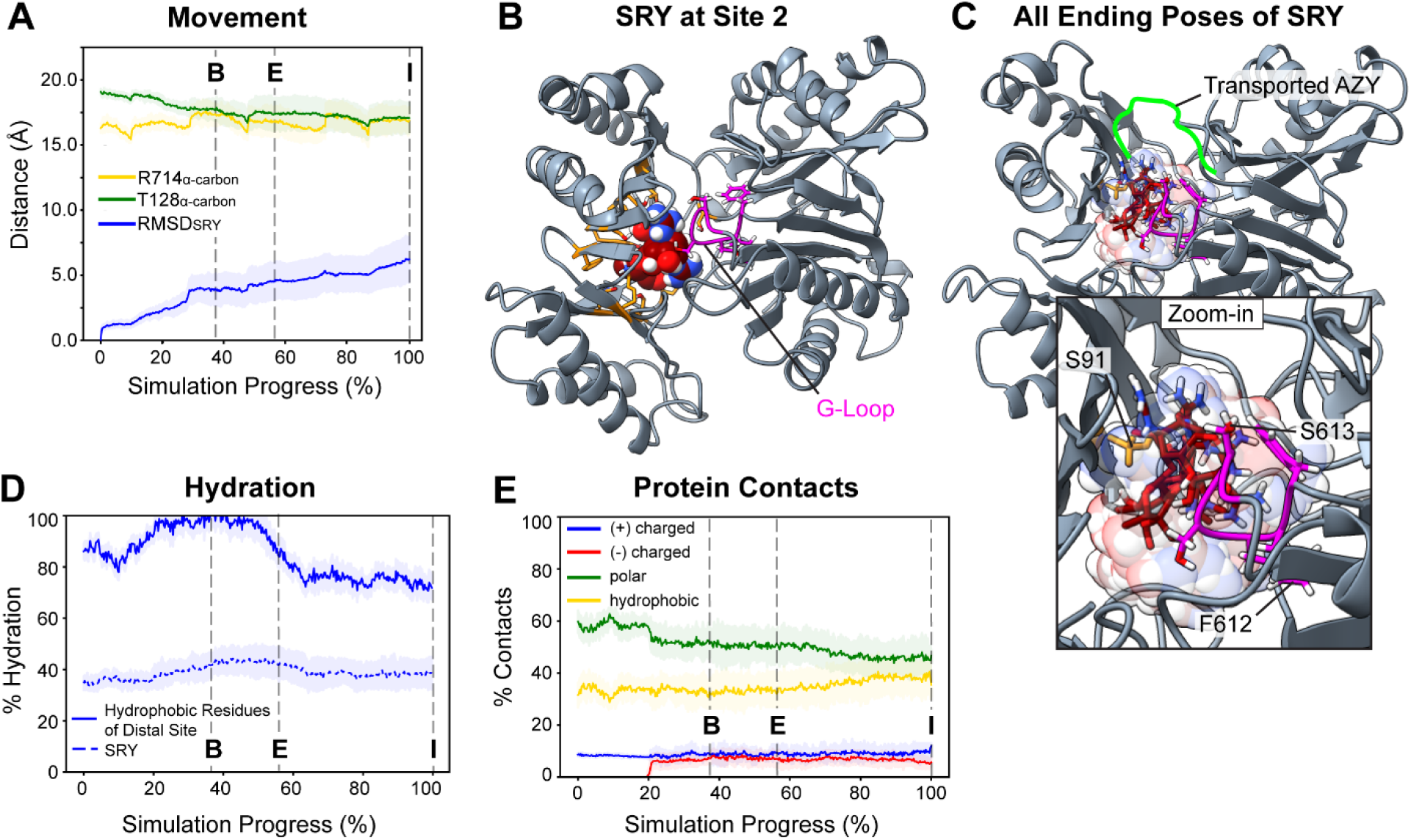
SRY is not Transported in Targeted MD Simulations from Site 2. **(A)** shows the RMSD_SRY_ from its starting position, the SRY_COM_ to T128α-carbon distance, and the SRY_COM_ to R714_α-carbon_ distance, over time. **(B)** shows SRY (maroon) at Site 1 in the Access Pocket. **(C)** shows all endpoints of SRY superimposed; one ending position of SRY, the position closest to fulfilling the criteria for “Transport”, is shown in opaque licorice representation. A green line marks the ending position of transported AZY for comparison. **(D)** shows the percent (%) hydration of the SASA_SRY_ or the SASA_distal hydrophobic_ over time. **(E)** shows the characteristics of MtrD-SRY contacts over time. In **(C)**, labeled residues in orange sticks interact with SRY in at least 80% of the TMD simulations. The G-Loop is shown in magenta.

### AZY Diffuses through the Periplasmic Cleft in an Unbiased Simulation

To avoid simulating multiple states of the PRN, we used Targeted MD techniques to model the conformational changes of MtrD^*16*^. However, Targeted MD is limited by the lack of control over simulation timescales (depending upon the Targeted MD implementation) and the use of biasing forces upon the protein. To address these limitations and complement our biased MD simulations, we performed an unbiased 1.5 μs simulation of AZY_neu_ at Site 2 using the AMBER pmemd-cuda MD engine (Figure 6A). AZY_neu_ was chosen because it exhibited the most frequent movement in Targeted MD Simulations. External biasing forces were not applied to the system, and the protonation states of the PRN were unaltered.

**Figure 6.**
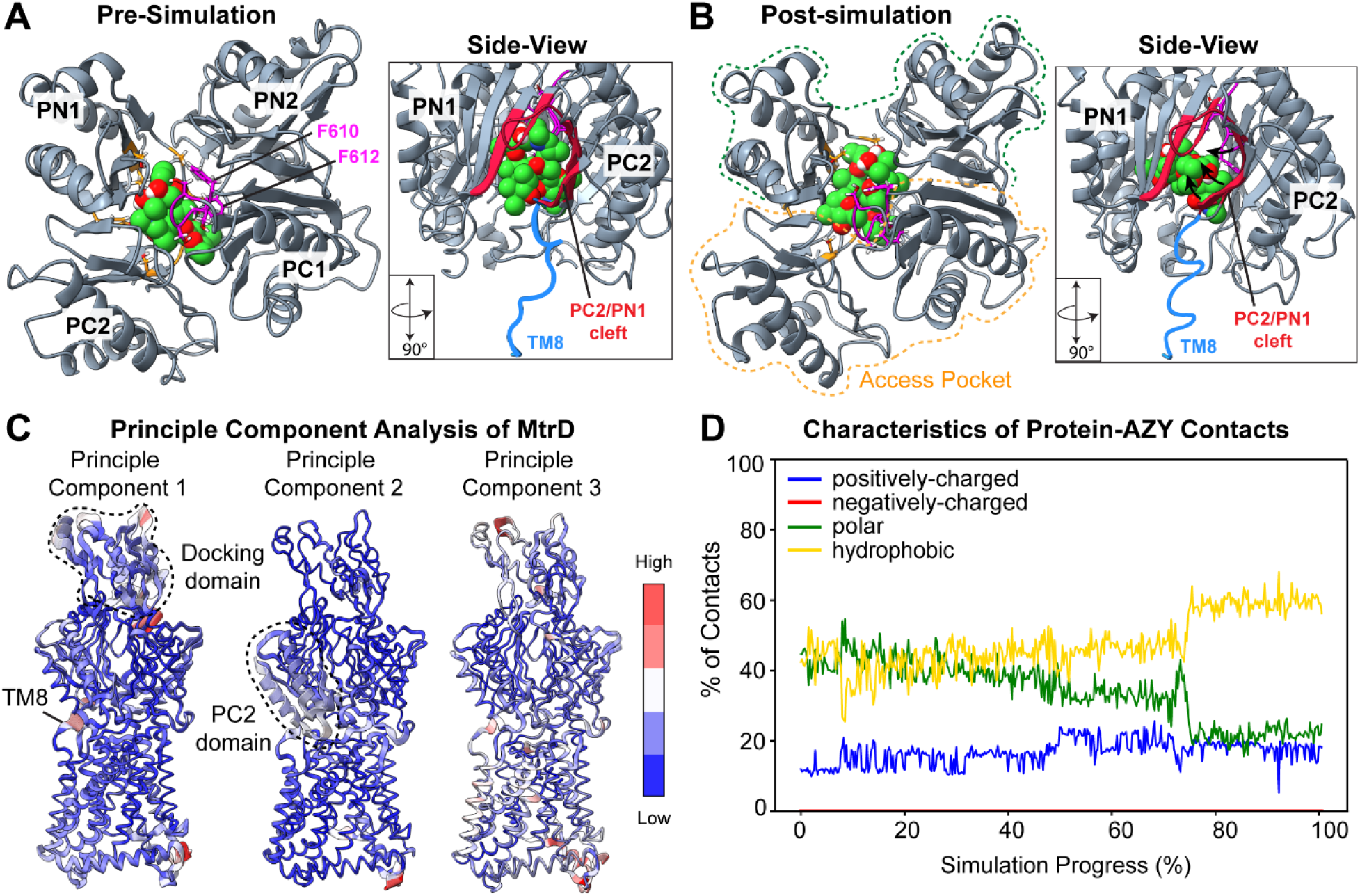
An Unbiased Simulation of AZY-Bound MtrD in the Access Conformation. **(A-B)** Pre- and post-simulation snapshots of AZY_neu_ and the periplasmic cleft, viewed from the top down, or viewed from the side (boxed view), showing AZY (green), the G-Loop (magenta), TM helix 8 (blue), and the PC2/PN1 cleft (outlined in red). Two phenylalanines of the G-Loop are labeled. AZY is in the starting position of Site 2. In **(B)**, arrows show how the PC2/PN1 cleft shifts. **(C)** Results of Principle Component Analysis on MtrD mapped onto the MtrD monomer. Red indicates areas of high structural fluctuation; blue indicates very low structural fluctuation. **(D)** The percentage of contacts between AZY and the periplasmic cleft that are charged, polar, or hydrophobic. **(E)** The hydration of the SASA of AZY or residues of the putative distal site throughout the simulation.

This unbiased simulation allowed us to address several outstanding questions. First, our Targeted MD simulations showed that AZY entered the Deep Pocket during the transition of MtrD from Access to Binding. If we observe movement of AZY into the Deep Pocket before the monomer has fully adopted the Binding structure, these data would not only correlate with our Targeted MD simulations, but also support the hypothesis that substrate movement into the Deep Pocket is governed by passive diffusion.

Second, MD studies of AcrB suggest that the transition from Binding to Extrusion requires energy from the PRN, and that the transition from Access to Binding does not^*11, 12, 17, 31*^. If this hypothesis is correct, we should see our AZY-bound Access monomer begin to transition to the Binding conformation. Structures of MtrDCR103 and AcrB show that TM8 adopts a more ordered structure, and PC2 opens wider (PDB IDs 6VKT, 5NC5)^*10, 19*^. Since the PRN remains unaltered, we would not expect MtrD to adopt the Extrusion structure. Whether we would observe MtrD fully adopt the Binding structure, and if any conformational changes would occur before AZY might enter the Deep Pocket in unbiased simulations, remain open questions.

In our unbiased simulation of AZY-bound MtrD in Access, AZY was indeed observed to diffuse from the proximal site into the Deep Pocket (Figure 6B, Table 1). Principle Component Analyses of MtrD revealed that structural fluctuations occurred in the PC2 domain, TM helix 8, and the docking domain (Figure 6C). Since MtrC and MtrE components were absent from our simulations, fluctuations in the docking domain are unsurprising. TM8 was observed to transition from a disordered loop to a more ordered structure, PC2 opened wider, and the PC2/PN1 cleft contracted (Figure 6A,B in blue, Movie S5). These changes are correlated with the transition from Access to Binding states. Notably, these conformational shifts began once AZY slipped fully into the Deep Pocket (Movie S5).

We found that MtrD-AZY contacts dominated by hydrophobic interactions at the end of the simulation (Figure 6D). Hydration of the SASA_AZY_ decreased from ~40% to ~35% over the course of the simulation; however, the SASA_distal hydrophobic_ was ≥ 70% hydrated for most the simulation. In combination with domain contact analyses, these data show that significant hydrophobic interactions between AZY and the distal site are not occurring, even in this unbiased simulation. The movement of AZY and the correlated motions of MtrD are shown in Movie S5. This unbiased simulation supports the hypothesis that substrate entry into the Deep Pocket, and the structural transition from Access to Binding, is correlated with a) the passive diffusion of a substrate into the Deep Pocket, and b) lack of an energy-requiring step from the PRN.

### The Molecular Landscape of the Periplasmic Cleft is Dynamic

In both Targeted and unbiased MD simulations, the MtrD substrate AZY was observed to move into the Deep Pocket during the transition from Access to Binding. Since both the Access and Deep Pockets are open in these states, substrate diffusion was occurring passively and *before* closure of the Access Pocket as the protein subsequently transitions to the Extrusion conformation. To investigate how the molecular characteristics of MtrD might facilitate substrate diffusion, we analyzed the Molecular Lipophilicity Potential (MLP) and the Electrostatic Potential (EP) of the periplasmic cleft at each stage of the putative transport cycle.

A previous analysis of MexB, which shares a 49% identity and similar substrate profile with MtrD, suggests that pump-ligand interactions involve substrate-complementary MLP/EP isosurfaces in the Access Pocket, whereas substrate movement involves weak dispersive forces contributed by mosaic isosurfaces^*33, 34*^. We found that the entrance of the Access Pocket of MtrD contains hydrophobic-amphiphilic complementary isosurfaces, and that the interior of the cleft consists primarily of mosaic isosurfaces. Additionally, we found that access to areas with specific MLP and EP signatures is alternately allowed and restricted throughout transport.

#### Molecular Lipophilicity Potential

MLP describes the 3D distribution of lipophilicity across a molecular surface, and is calculated by summing the lipophilic contributions of molecular fragments upon the surrounding environment^*33*^. A positive (+) MLP value indicates a *lipo*philic region (Figure 7, gold regions), and a negative (-) MLP indicates a *hydro*philic region (Figure 7, teal regions). MtrD transports diverse hydrophobic or amphipathic molecules^*8*^. When MtrD is in Access, PC2 and PN1 form mosaic neutral/hydrophilic isosurfaces (Figure 7A, left). On the opposite wall of the cleft, PC1 and PN2 provide lipophilic isosurfaces similar to those observed in MexB (Figure 7A)^*34*^. In Binding, PC2 and PN1 form neutral/hydrophilic mosaic isosurfaces (Figure 7B, left image). PC1 and PN2 appear to create a ‘lipophilic highway’ that extends directly through the cleft (Figure 7B, right image), but the G-Loop prevents consistent contact with PC1 and PN2 as substrates move through the cleft^*8*^. As shown by our docking studies, the predicted binding modes of AZY curved to the left of the G-Loop, contacting PC2 and PN1 (Figure S2). In Extrusion, access to PC1 and PC2 is restricted by the closure of the Access Pocket, and the available MLP isosurfaces to a ligand are hydrophilic/neutral mosaics, apart from the lipophilic “plug” formed by the G-Loop (Figure 7C). In Intermediate, the Deep Pocket constricts and the Access Pocket remains closed (Figure 7D).

**Figure 7.**
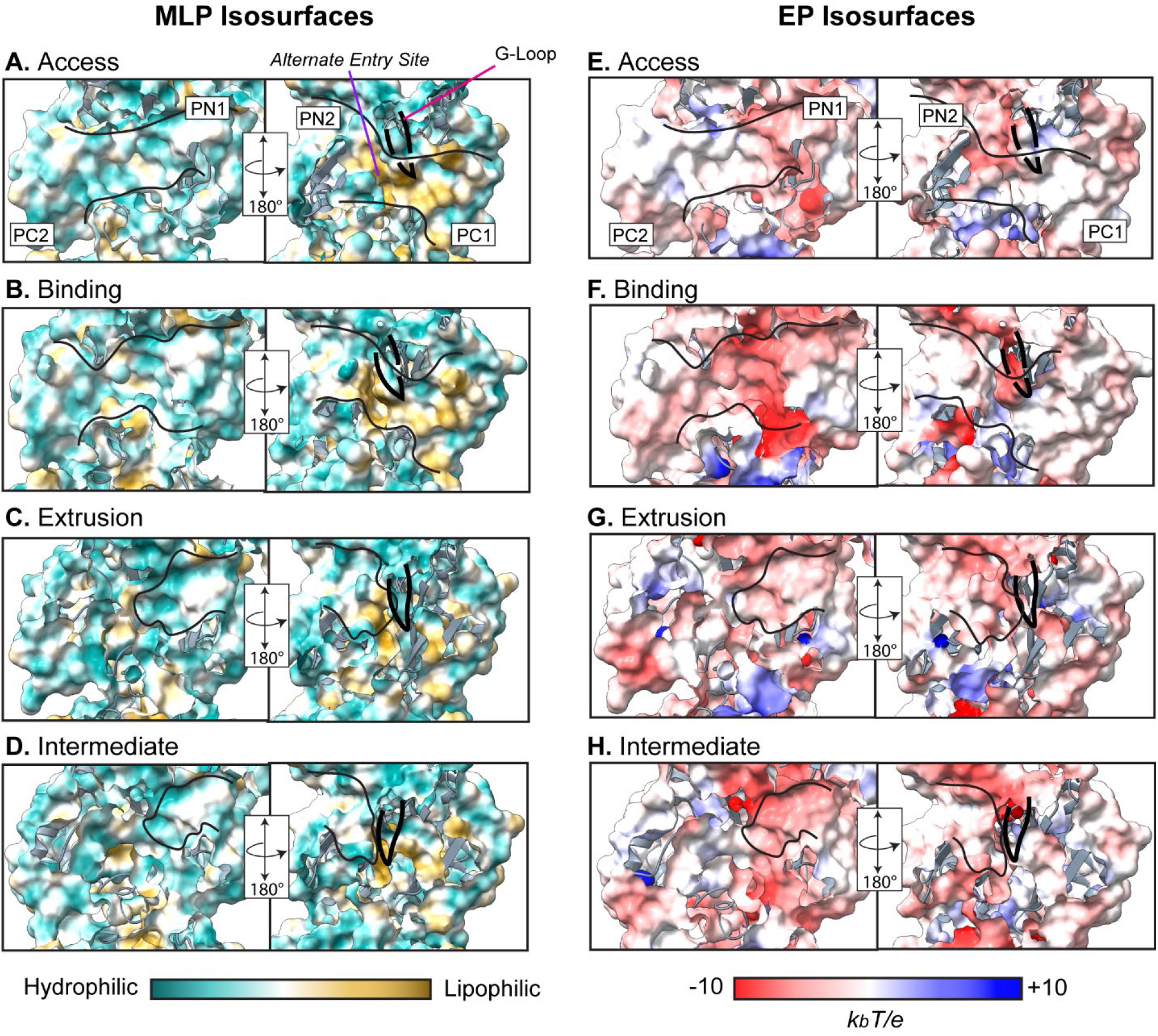
The Dynamic MLP and EP Isosurfaces of the Periplasmic Cleft of MtrD. Panels. (**A-D)** show the Molecular Lipophilicity Potential (MLP) isosurfaces plotted on the molecular surface representation of the periplasmic cleft with MtrD in **(A)** Access, **(B)** Binding, **(C)** Extrusion, and **(D)** Intermediate. MLP is colored teal to gold; teal regions are more hydrophilic, and gold regions are more lipophilic. **Panels (E-H)** show the Electrostatic Potential (EP) isosurfaces of the periplasmic cleft with MtrD in **(E)** Access, **(F)** Binding, **(G)** Extrusion, and **(H)** Intermediate. EP is colored red to blue, from negative (−10 kbT/e) to positive (+10 kbT/e) potential, where kb is the Boltzmann constant, T is the absolute temperature (310K), and e is the electron charge. The available conformational space within the cleft is approximated by black lines. The approximate position of the G-Loop is marked in each structure by a black line. MLP is colored teal to gold, from hydrophilic to lipophilic. EP is colored red to blue, from negative to positive. MLP was calculated using ChimeraX^*35*^, EP was calculated using PDB2PQR and the APBS server^*36*^.

#### Electrostatic potential (EP) isosurfaces of the periplasmic cleft

Access to electrostatic potential (EP) isosurfaces of the periplasmic cleft changed dynamically throughout the transport process. Figure 7E-H show the EP of MtrD. In Access, the PC1 and PC2 domains consist of mostly neutral EP isosurfaces (Figure 7E). In Binding, a negative patch was observed to form near the interface of PN1 and PC2, in a region located behind the G-Loop (Figure 7F). The opposite wall of the cleft (formed by PC1 and PN2) is mostly neutral, with the G-Loop contributing negative electrostatic interactions. In the Extrusion and Intermediate conformations, the Deep Pocket is the only available conformational space to a bound ligand. PN2 consists of neutral/negative mosaic EP surfaces, whereas PN1 contributes more negative isosurfaces (Figure 7G-H). Thus, the electrostatic environment within the periplasmic cleft of MtrD is largely negative-neutral throughout the putative efflux cycle. Many MtrD substrates – like AZY_h1_ – are amphiphilic or weakly positively charged; therefore, the EP isosurfaces of the periplasmic cleft are complementary to the substrate profile of MtrD.

## Discussion

Using Targeted and unbiased MD simulations, we observed the simulated transport of AZY by MtrD without pulling (or pushing) upon the ligand itself. Including a fourth “Intermediate/Resting” structure^*16*^ increases the transport distance of AZY, suggesting that this state might be a “re-set” at the end of the efflux cycle. AZY was observed to take an unexpected pathway along PN1 and bypassed the distal site altogether. In Targeted MD simulations, we also observed the rejection – or retention – of SRY from its initial starting positions within the Access pocket, confirming that SRY is not a transport substrate nor is it likely an effective inhibitor of MtrD^*6*^. Through analyses of the MLP and EP of the periplasmic cleft, we found that access to specific isosurfaces was alternately restricted and allowed throughout transport.

Taken together, our data suggest that multiple transport pathways through the periplasmic cleft may exist – even for large macrolides like AZY. Our data also support the hypothesis that substrate diffusion through the Access Pocket and into the Deep Pocket occurs slowly, and in the absence of changes in the Proton Relay Network^*11*^. We hypothesize that the transport of AZY by MtrD is mediated through a combination of 1) alternating access to specific isosurfaces of the periplasmic cleft, 2) a peristaltic-like squeezing caused by the sequence of conformational changes that MtrD undergoes, and 3) hydration of the substrate throughout transport.

### Dynamic Molecular Characteristics of the Periplasmic Cleft

Through analyses of the MLP and EP of the periplasmic cleft, we found that ligand access to hydrophobic areas of the pump was alternately opened and restricted by the closure of the Access Pocket. These analyses also revealed that the periplasmic cleft of MtrD contains 1) hydrophobe-amphiphile-complementary isosurfaces at the entrances, and 2) mosaic isosurfaces along the interior. Through their analysis of the MtrD homologues MexB and AcrB, Ramaswamy et al. and Vargiu et al. suggested that mosaic isosurfaces – along with the presence of water – would facilitate the smooth diffusion of substrates through the periplasmic cleft^*31, 34*^. The results of both our Targeted and unbiased MD simulations support this hypothesis. Without applying forces to the ligand itself, AZY was able to diffuse along these mosaic isosurfaces and towards the funnel domain. We also found that AZY was consistently hydrated throughout the transport process, suggesting that water may indeed play a role in substrate movement through the cleft^*31*^. We note that mosaic isosurfaces are advantageous for transport, as they may prevent substrates from stabilizing – or interacting too strongly – with the cleft interior.

### AZY did not Exit into the Funnel Domain

We did not observe the movement of AZY into the funnel domain in our MD simulations. This is unsurprising, as an earlier MD study of AcrB and its substrate Doxorubicin relied upon steered MD forces to pull Doxorubicin into the funnel domain, thereby necessitating dissociation^*31*^. However, with the inclusion of the putative Intermediate state in our target sequence, AZY was observed to end in (2.5 - 3 Å increase in distance traveled) a better position for release/dissociation (Figure 4D, ‘Intermediate’). To our knowledge, no computational simulation of an RND transporter has modeled substrate release into the funnel/docking domain without the use of biasing forces upon the substrate itself^*8, 37*^. These data suggest that substrate release may occur on a longer timescale than is currently reasonable to simulate.

### The Transition from Access to Binding may not Require Changes in the PRN

Our results are in line with MD studies of the MtrD homologue AcrB, which suggest that the transition from Binding to Extrusion is the energy dependent phase^*31*^. Our results are also supported by the MD study of MtrD performed by Chitsaz et al., in which progesterone was observed to spontaneously move into Access Pocket, past the G-Loop, and into the Deep Pocket – all in the absence of biasing forces, and without changes to the Proton Relay Network of MtrD^*8*^. Chitsaz et al. observed the movement of progesterone through the periplasmic cleft in ~40 ns; we postulate that the increased movement speed (relative to our simulation of AZY) could be due to the significant discrepancy in size between progesterone and AZY (~315 Da for progesterone vs. ~749 Da for AZY). Their simulations began with MtrD in the Access conformation, again suggesting that substrate movement into the Deep Pocket occurs before the monomer fully adopts the Binding structure, and occurs through passive diffusion.

### The “PN1 Route” and the Distal Site Re-examined

Simulations of ligand-bound RND pumps have relied upon biasing forces to pull transport ligands through – or out of – the periplasmic cleft^*31, 37*^. While these simulations can be quite informative, these techniques also bias the path of the ligand towards a user-determined vector. In the work presented here, we sought to simulate substrate transport by MtrD without pulling upon the transport ligand itself. Through these simulations and found that AZY took an unexpected pathway – the “PN1 Route” (Figure 8A, orange arrow). We observed that this alternate route bypassed the distal site and was mediated by water, PN1 and the G-Loop. These results conflict with co-crystallization complexes of erythromycin (ERY) bound MtrDCR103. This structure (6VKT) shows ERY bound at the distal site, with the MtrD monomer in the Binding conformation. At no point in any of our simulations was AZY bound in a conformation like that of erythromycin in the Deep Pocket of MtrDCR103. Two questions arise – 1) is the “PN1 Route” an artifact of the TMD simulations, and 2) are there multiple transport pathways through the periplasmic cleft?

**Figure 8.**
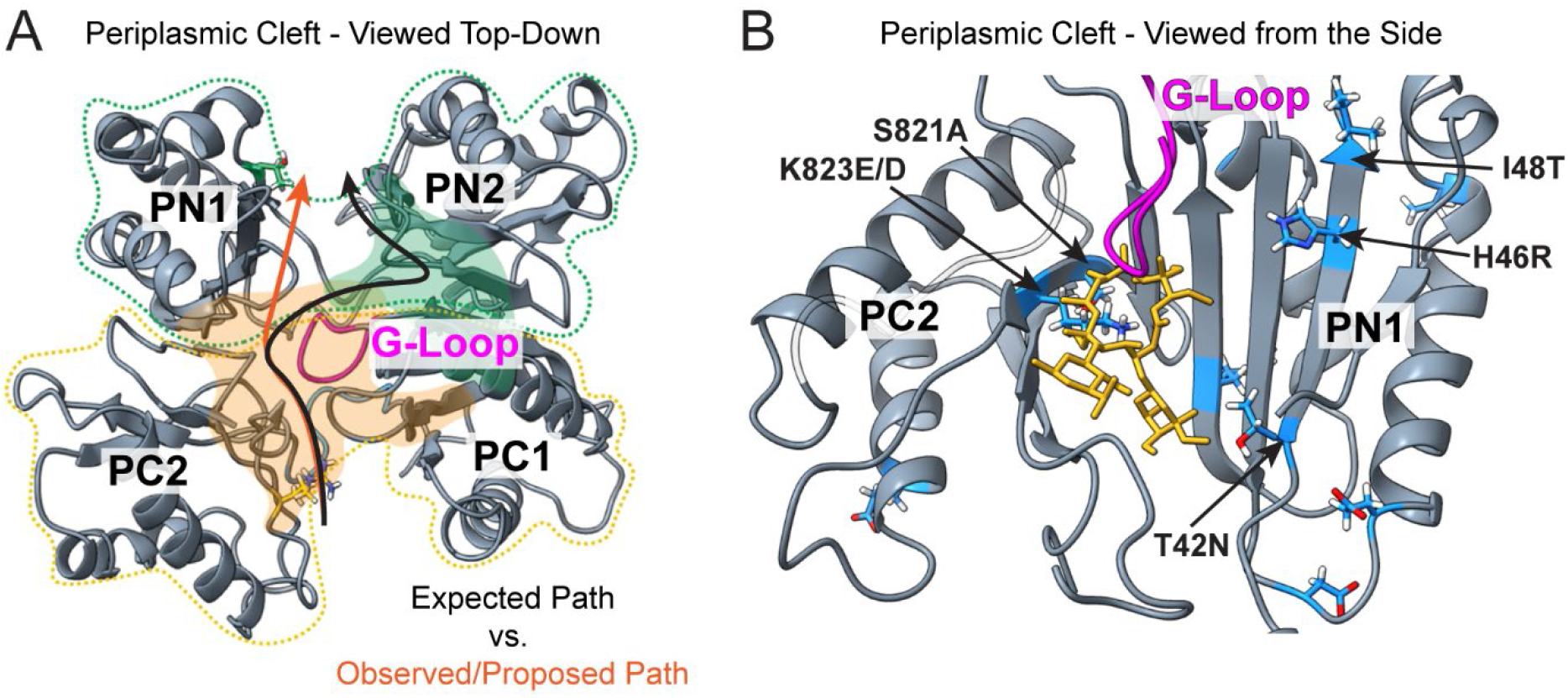
PC2 and PN1 Mutations Confer AZY Resistance in *N. gonorrhoeae*. Here we show two views of the periplasmic cleft. **(A)** shows the periplasmic cleft viewed from the top down. A black arrow shows the expected path of AZY during transport, based upon co-crystallization of ERY at the Distal Site (green shaded area, composed of residues from PC1 and PN2)^*10*^. An orange arrow shows the observed path of AZY during transport, dubbed the “PN1 Route”. **(B)** shows the periplasmic cleft viewed from the side, with MtrD domains labeled. AZY is shown in yellow licorice representation. The G-Loop is in magenta. Residues in blue sticks are mutations found in the AZY-resistant GCGS0276 strain of N. gonorrhoeae^*38*^. Transforming the non-resistant 28BI strain with the PC2 and PC1 domains of GCGS0276 increased the AZY MIC of 28BI fourfold^*38*^. 28BI transformants with other domains of GCGS0276 did not exhibit an increase in the MIC of AZY.

We found that a recent study by Wadsworth et al. supports the existence of the “PN1 Route” for AZY^*38*^. Wadsworth et al. showed that mutations in the PC2 and PN1 domains of MtrD correlate with increased resistance to AZY in clinically obtained strains of *N. gonorrhoeae*. In a particularly elegant experiment, Wadsworth et al. transformed the *mtrD* locus in a non-AZY-resistant strain (*N. gonorrhoeae*, 28BI) with two portions of the *mtrD* locus from an AZY-resistant strain (*N. gonorrhoeae*, GCGS0276). These two regions produce the PN1 and PC2 domains of MtrD and contain mutations in areas that might contact AZY during transport (20 mutations total) (Figure 8B, blue residues). The substitution of these two domains from the AZY-resistant strain increased the (28BI, non-resistant strain) MIC for AZY fourfold^*38*^. Importantly, 28BI transformants involving other regions of MtrD from GCGS0276 did not raise the 28BI MIC of AZY, suggesting that PN1 and PC2 are indeed involved in the efflux of AZY^*38*^. These experiments strongly support the hypothesis that PN1, and by extension PC2, is involved in transport of AZY, and correlate well with the results of our TMD experiments.

How do we reconcile these results with prior studies of the distal site? Indeed, mutations in the distal site of MtrD (F136A, F176A, and F623C) correlate with a decrease in the MIC (Minimum Inhibitory Concentration) of the antibiotics rifampin (~823 Da), novobiocin (~613 Da), and oxacillin (~401 Da)^*8*^. None of these antibiotics are macrolides like AZY, but they are considered “bulky” substrates, thus a decrease in the MIC indicates that mutations in the distal site are somehow disrupting the transport of these antibiotics. Indeed, studies of AcrB have shown that conformational flexibility in the G-Loop is critical for the transport of large substrates^*15, 39*^. Specifically, through MD simulations of AcrB with an inhibitor bound at the distal site, Vargiu et al. show that these distal site-bound inhibitors may interact with the G-loop, likely reducing its flexibility, which would impair transport of large substrates^*39*^. Thus, in addition to providing an alternate entrance for small substrates, the distal site may contribute to transport by mediating the conformational changes of the G-Loop.

An additional important question that remains is whether the conformational change from Binding to Extrusion is enough to extrude a large substrate out of the distal site and into the funnel domain. To investigate, we modeled the structural transition from Binding to Extrusion using the erythromycin-bound monomer of MtrD_CR103_ and the Morph capability of ChimeraX (6VKT, Movie S6). We found that the distal site constricts significantly during this structural transition. While this constriction may be sufficient to squeeze a substrate up and into position for dissociation, it is unclear whether enough force is provided to extrude the substrate. It appears, however, that conformational changes of the distal site are necessary during the transition from Binding to Extrusion. Thus, the disruption of this site might hinder this necessary step in substrate transport.

Therefore, the role of the distal site in macrolide transport remains unclear, and more studies are warranted. We suggest that multiple transport pathways through the cleft – may exist, even for large macrolide antibiotics like AZY. The originally proposed transport pathway incorporates close interactions with the distal site behind the G-Loop, which is formed by the PN2 and PC1 domains (Figure 8A, black arrow, distal site is shaded green). The alternate transport pathway bypasses these distal site interactions, with the substrate instead traveling along the PN1 domain (Figure 8A, orange arrow).

### Multiple transport pathways through the periplasmic cleft

In conclusion, our data suggest that substrate transport through the periplasmic cleft of MtrD depends upon a combination of diffusion and peristaltic motions of the periplasmic cleft, resulting in gated access to areas with variable charge and lipophilicity isosurfaces. We have identified a low energy, peristalsis-complementary diffusion path for AZY through the periplasmic cleft that does not involve interactions with the putative distal site^*17*^. Our results suggest that multiple pathways, or residue contact pathways, may exist within the periplasmic cleft for substrates of MtrD.

We suggest the following transport mechanism for macrolides by MtrD. (1) A macrolide associates with and passes the selectivity filter formed by the Access Pocket. (2) The macrolide diffuses slowly through the periplasmic cleft, contacting residues of the proximal site, and eventually associating with the G-Loop^*10*^. (3) As the macrolide slips past the G-Loop and into the Deep Pocket, the MtrD monomer undergoes the transition from Access to Binding, opening the periplasmic cleft even wider. (4) Once in the Deep Pocket, the macrolide interacts with the distal site or with the PN1 domain, and close contacts with the G-Loop are maintained. (5) Changes in the Proton Relay Network power the pump’s transition from Binding to Extrusion, resulting in the closure of the Access Pocket and squeezing the substrate towards the funnel domain^*12*^. (6) The monomer reaches the Extrusion conformation and “waits” for substrate release. (7) The monomer might relax and adopt the Intermediate structure during its transition back to Access, as the Intermediate structure mostly involves a peristaltic constriction of the cleft.

## Acknowledgements

The authors would like to acknowledge and thank Dr. James W. McCormick for his valuable and constructive suggestions and guidance during the planning and development of this work. We would also like to thank Amit Kumar for his continual support and patience with our use of the *Maneframe II* computer.

The authors declare no competing financial interest.

## Materials and Methods

### Software

Missing residues in the 4MT1 structure of MtrD were repaired with Modeller v. 9.24 (residues 1, 494-507, 671-672, 1041-1056)^*5, 24*^. The complete MtrD homotrimer was built in VMD (Visual Molecular Dynamics, v 1.9.4) using the crystallographic coordinates in the 4MT1 PDB file^*40*^. A heterogeneous bilayer consisting of POPE (70%), POPG (20%), and cardiolipin (10%) was created using the CHARMMGUI Membrane Builder with CHARMM36 topology^*41–43*^. This heterogenous bilayer was then minimized, heated to 310K, and equilibrated using the NAMD inputs created by CHARMMGUI; settings were maintained at default, with the exception of the simulation temperature, which was changed from the default of 298K to 310K. The repaired MtrD homotrimer was minimized to relax the modeled loops, and then embedded into the equilibrated heterogeneous membrane using coordinates from the Orientation of Proteins in Membranes (OPM) Michigan database for 4MT1^*44*^. Lipids were retained in the central pore of the protein. The protein-membrane system was solvated with TIP3 water and ionized to a concentration of 0.15M NaCl using the Solvate and Autoionize plugins of VMD. The solvated and ionized protein-membrane system consisted of ~339,000 atoms, and was equilibrated for 100 ns prior to use for further experiments.

All Molecular Dynamics (MD) simulations were performed with NAMD 2.12 using CHARMM36 force fields and topology unless otherwise stated. Targeted Molecular Dynamics simulations were performed using NAMD 2.12 and in-house scripts written in *Tcl*. The 1.5 μs simulation of MtrD and AZY was performed with AMBER18 using the ff14SB, Lipid17, GAFF2 and TIP3p force fields employing the pmemd.cuda-DPFP molecular dynamics engine^*45–47*^. The CHARMM-GUI Ligand Reader and Modeller was used to format and parameterize ligands for simulations^*41, 42*^. PROPKA3.1 and the Henderson-Hasselbalch Equation was used to estimate ligand protonation states at a pH of 7.4. *Autodock Vina 1.1.2* was used to dock AZY or SRY ligands to the drug binding domain of MtrD. Autodock Tools was used to define dock boxes^*23, 48*^. Bio3D was used for analysis and generation of target structures for Targeted MD simulations^*49*^. Protein images were generated using UCSF ChimeraX^*35*^. Data analysis was performed using in-house scripts written in *Tcl*, R, and python. Figures were prepared with Adobe Illustrator. Movies were prepared with ChimeraX, VMD, and Adobe Procreate.

### Ligand docking to the drug binding pocket of MtrD

To generate a starting position for our ligands of interest, we docked azithromycin and streptomycin with the periplasmic cleft of wild-type MtrD. The structures of the MtrD substrate azithromycin (AZY) and the non-substrate streptomycin (SRY) were downloaded from PubChem and converted to 3D structures using OpenBabel v. 2.3.2^*50*^. The full-length (repaired) MtrD monomer and ligands were converted to PDBQT files using AutoDock Tools v. 1.5.6^*48*^. AutoDock Vina 1.1.2 was then used to dock each ligand with four overlapping boxes that encompassed the entire periplasmic cleft of MtrD (Figure S2)^*23*^. The docking exhaustiveness parameter was set to 128 replicates to ensure reasonable coverage of the docking regions; default exhaustiveness for Autodock Vina is 8. Docking results were processed using in-house bash scripts, producing the top 9 poses of each ligand per dock box, ranked by binding affinity (Figure S2).

To select ligand start sites for MD simulations, the resultant docking poses were then processed using AutoDock Tools. The purpose of these docking experiments was to generate a plausible starting position for the ligand within the periplasmic cleft, not to evaluate individual estimated binding affinities. Vina has an estimated standard error in calculating binding energies for small molecule redocking experiments of 2.85 kcal/mol^*23*^. Similar experiments for calculating the standard errors of affinity estimates for peptide- or protein-ligand complexes is expected to be much higher, due in large part to the significantly greater conformational degrees of freedom allowed for the ligand of interest. Consequently, the top 9 docking poses were evaluated according to position within the periplasmic cleft, and not to individual estimated affinities.

We found that the overlapping dock boxes produced some identical docking poses (Figure S2). From each cluster of identical poses, one representative pose was selected randomly. After the elimination of identical poses, three poses were selected for both AZY and SRY. For MD simulations with each ligand, we selected one docking pose at the Mid-Point of the cleft but within the proximal binding site – this became Start Site 1. We chose a second pose at the entrance of the periplasmic cleft – this became Start Site 2. The PDBQT files of ligands in each selected pose were converted to PDB format and “all atom” representations using Open Babel 2.3.2, since the PDB to PDBQT processing (for docking) removes all non-polar hydrogens^*50*^.

The CHARMMGUI Ligand reader and Modeler were then used format the ligands as PDBs, to create parameter files for simulations with NAMD or AMBER, to create various protonation states if applicable, and to check for stereochemistry issues^*41, 42*^. Using available data (if possible) and analyses with Propka3.1 and OpenBabel, ligand protonation states were assessed at a physiologically relevant pH of 7.4^*27, 50*^. At this pH, according to our calculations, SRY is a triple cation with only ~0.02% of molecules being double cations at pH 7.4. Therefore, we simulated the completely protonated form of streptomycin (SRY). For azithromycin (AZY), there are two ionizable sites (Figure 2). For the first ionizable site, the pKa is ~8.5-8.74, meaning that ~4.6-7.9% of AZY are unprotonated at a pH of 7.4^*25, 28*^. For the second site, the pKa is 9.45, meaning that ~1% of AZY are unprotonated at this site^*28*^. Since it is thought that the neutral, unprotonated form of AZY may more readily diffuse through cell membranes, we included all three protonation states of AZY in our simulations.

We note also that pKa values may change depending upon the surrounding environment, and that the pKa of amines – of which there are two present on AZY – is expected to increase when moving from an aqueous to a more hydrophobic environment. This may result in higher percentages of the unprotonated form of these AZY in more hydrophobic environments. While these environments may increase the proportion of the unprotonated species of AZY, they are not expected to significantly change the protonation state of SRY (Figure 2).

### TMD simulations of the ligand free system with NAMD

TMD simulations, and the unbiased relaxation MD simulations that preceded TMD, were performed using NAMD 2.12 with the CHARMM36 forcefield, a timestep of 2 fs, and a non-bonding cutoff of 12 angstroms^*43, 51*^. Simulations were performed with constant temperature and pressure (NPT) in a periodic cell using Langevin temperature and pressure control, and Particle Mesh Ewald electrostatics. The temperature was maintained at 310K. As preparation for subsequent docking experiments and TMD simulations, the ligand-free system was minimized, heated to 310K, and equilibrated for 100 ns. An equilibrated MtrD monomer was extracted from the end of the final simulation and used for docking ligands and for aligning target structures for TMD simulations.

#### Preparing the protein-ligand system

After docking, each ligand was converted to a PDB and then uploaded, checked for structural or conformational issues, and parameterized using the CHARMM-GUI Ligand Reader and Modeller. After adding the ligand into the system and removing any overlapping water molecules using VMD, the new system was minimized, heated to 310K, and equilibrated in unbiased MD simulations for 50 ns. Using a short TMD simulation, the alpha carbons of the equilibrated system were then guided to the starting coordinates of the 4MT1 crystal trimer.

#### Targeted Molecular Dynamics Simulations with NAMD

To mimic a putative drug transport cycle of MtrD, TMD simulations were performed as previously described^*52, 53*^ using target coordinates derived from structures of MtrD homologue(s) (Figure S1). In our TMD simulations, alpha carbons of the MtrD backbone are guided to an RMSD ≤ 0.7 Å from the target coordinates. Forces were not applied to the ligand of interest, and protein sidechains move freely. When simulations were performed, no published structures of MtrD in various conformations were available, so we used structures derived from AcrB from *E. coli*, which shares a sequence identity of 48.6%, and CmeB from *C. jejuni*, which shares a sequence identity of 38.07%, with MtrD. Structures used for TMD simulations were the 4DX5 (1.9 Å) and 5NC5 (3.2 Å) structures of AcrB, and the 5LQ3 (3.5 Å) structure of CmeB (Figure S1). A comparison of the normal (“wildtype”) MtrD, MtrDCR103, and the structures used in our TMD simulations, is provided in Figure S6. Using the structurally homologous Cα atoms of MtrD for each structure, target structures were superposed with the equilibrated MtrD-ligand system using an in-house script and the Bio3D module of R.

In subsequent TMD simulations on equilibrated protein-ligand systems, forces were applied using in-house *tcl* scripts in NAMD^*53*^. These forces were applied to structurally homologous Cα atoms of the ligand-bound protomer to gently guide the Cα atoms toward the respective target coordinates. The magnitudes of these forces were inversely proportional to the RMSD (root mean squared deviation) of the distances separating the selected Cα atoms from their target coordinates. Cα atoms were pushed to RMSD values of ≤ 0.7 Å from their target coordinates. Significant steric clashes were not observed between protomers, even though only one protomer was guided through conformational changes. These short timescale TMD simulations, on average, ran for approximately 2 nanoseconds each. While these TMD simulation timescales are quite short, they are nonetheless useful^*20, 52*^, as they allow us to test if the peristaltic motions of the periplasmic cleft can indeed result in the movement of a ligand through the cleft. These TMD experiments thus provide the basis for longer timescale experiments.

Analysis of the protein-ligand interactions for each system were performed using *tcl* scripts in VMD and scripts written using the Bio3D R module. At equal intervals throughout each simulation, the number and characteristics of ligand-protein contacts were determined. Results are reported either for individual simulations or the averages for all twenty simulations per ligand, per starting site.

We simulated three protonation states of the MtrD substrate azithromycin (AZY): AZY_neu_, a neutral, unprotonated form of azithromycin; AZY_h1_, a singly protonated, positively charged form of azithromycin; and AZY_h2_, doubly protonated, positively charged form of azithromycin (Figure 2). As a negative control, we tested streptomycin (SRY), a known non-substrate of MtrD, and a triple cation at physiological pH. Two start sites were tested for each ligand −Site 1, in which the ligand was located in between the AP and the DP near the G-Loop, and was associating with the proximal binding site, and Site 2, in which the ligand was at the entrance of the periplasmic cleft and was associating with the foremost residues of the proximal binding site (Figure 2).

The center of mass coordinates, and ligand RMSDs from starting positions, were calculated using Tcl scripts in VMD. Protein-ligand systems were oriented such that the membrane is in the X – Y coordinate plane; therefore, *positive* vectorial movement along the X axis indicates movement through the periplasmic cleft *towards* the central region of MtrD, as would be expected during ligand transport. In contrast, *negative* vectorial movement along the X axis indicates movement *away* from the central region and towards the periplasm. Both R714 and T128 were chosen as points of reference based upon their positions within the cleft. Additionally, both residues primarily move on an axis perpendicular to the direction of transport, and exhibit an overall RMSD of ~2.5 Å. Since biased external forces were not applied to the ligand, any movement through the protein should only be dependent upon the conformational changes of the MtrD protein and the limited diffusional possibilities of the transport substrate, which are dependent on the protein conformations.

#### A long timescale MD simulation of AZY-bound MtrD

Using the AMBER pmemd-cuda MD engine, we performed a GPU-accelerated, 1.5 μs simulation of the MtrD homotrimer with AZYneu bound at Site 1 in the periplasmic cleft^*46*^. The ligand was parameterized with AMBER antechamber using the GAFF2 force field, and the protein-ligand system was built in AMBER tLEAP^*46, 54, 55*^. Except for parameterization files, the system was identical in composition and size to the system run with NAMD, except that the membrane did not contain cardiolipin; the heterogeneous membrane (POPE (70%) and POPG (30%) only), water, and ions were maintained. The system was first relaxed in unbiased equilibration MD simulations for 200ns, and then allowed to run freely for a total of 1.5 μs. Analysis was performed using AMBER cpptraj, UCSF ChimeraX, the PDB2PQR server, in-house Tcl scripts with VMD or with python, and with Bio3D in R^*35, 36, 40, 46, 49, 56*^.

### Molecular Lipophilicity Potential

The molecular lipophilicity potential (MLP) describes the three-dimensional distribution of lipophilicity across the molecular surface of a molecule or protein. The MLP at a point in space (k) can be calculated using the following equation^*33*^, where N is the number of molecular fragments, Fi is the lipophilic contribution of each molecular fragment (i), and the distance function f(d_ik_) describing the distance between the point (k) to the molecular fragment *i*:

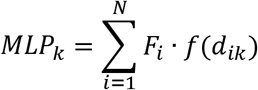

The sum of all MLP values for the molecular surfaces of the periplasmic cleft yields the Lipophilic Index (LI) of that region, defined as:

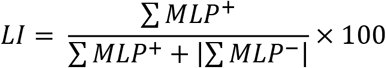

MLP^+^ denotes regions with a positive, or lipophilic, MLP value; MLP^-^ denotes regions with a negative, or hydrophilic, MLP value. If the fragmental contributions of the MLP^+^ and the MLP^-^ of a region sum to roughly zero, then the region is classified as MLP “neutral”. MLP calculations were performed and visualized using ChimeraX^*35*^.

### Electrostatic Potential

The electrostatic potential (EP) surfaces of the periplasmic cleft were calculated using the APBS/PDB2PQR server and visualized using ChimeraX^*36, 57, 58*^. All EP calculations were performed at 310K with all other Poisson-Boltzmann parameters at default. EP calculations were performed on MtrD both in presence and absence of AZY by isolating PDB “snapshots” of the ligand-free MtrD monomer from specific timepoints in the simulation; subsequent EP calculations were performed on these PDB snapshots using the APBS/PDB2PQR server.

### Percent hydration of solvent accessible surface area (SASA)

The percent (%) hydration of the available SASA of each ligand was calculated as follows:

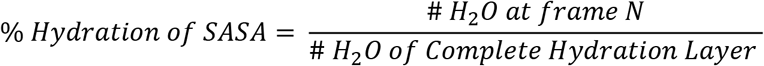

With N being the individual frame, or timepoint, analyzed. The complete potential hydration layer describes the number of water molecules that would surround the equilibrated ligand if it was immersed freely in solvent. The complete potential hydration layer was calculated by 1) immersing each ligand into a water box, 2) neutralizing the system with 0.15 mol/L NaCl, 3) minimizing the solvated system and heating it to 310K, and 4) simulating the ligand with free MD for 20 ns. The position of water molecules within three different radii of AZY or SRY (radii of 3 Å, 4 Å, 5 Å, and 6 Å) were assessed to determine which radius describes a complete hydration layer around the ligand of interest. For both AZY and SRY, the optimal radius was water within 4 Å of any atom of the ligand. The number of water molecules to completely hydrate each ligand in its free-MD relaxed state was determined to be as follows: AZY_neu_, 107 water molecules; AZY_h1_, 103 water molecules; AZY_h2_, 104 water molecules; SRY, 83 water molecules. The SASA was calculated using the “measure” function of VMD.

## Notes

### Competing Interest Statement

The authors have declared no competing interest.

### Summary of Updates

The text has been significantly edited for brevity and clarity. We have added to our analyses as well, and honed the focus of the paper.

## References

[1] Wade, J. J., and Graver, M. A. (2003) Survival of six auxotypes of Neisseria gonorrhoeae in transport media, J Clin Microbiol 41, 1720–1721.

[2] Rowley, J., Vander Hoorn, S., Korenromp, E., Low, N., Unemo, M., Abu-Raddad, L. J., Chico, R. M., Smolak, A., Newman, L., Gottlieb, S., Thwin, S. S., Broutet, N., and Taylor, M. M. (2019) Chlamydia, gonorrhoea, trichomoniasis and syphilis: global prevalence and incidence estimates, 2016, Bull World Health Organ 97, 548–562P.

[3] St. Cyr S, B. L., Workowski KA, et al. (2020) Update to CDC’s Treatment Guidelines for Gonococcal Infection, 2020, In MMWR Morb Mortal Wkly Rep 2020, pp 1911–1916.

[4] Nikaido, H. (2018) RND transporters in the living world, Res Microbiol 169, 363–371.

[5] Bolla, J. R., Su, C. C., Do, S. V., Radhakrishnan, A., Kumar, N., Long, F., Chou, T. H., Delmar, J. A., Lei, H. T., Rajashankar, K. R., Shafer, W. M., and Yu, E. W. (2014) Crystal structure of the Neisseria gonorrhoeae MtrD inner membrane multidrug efflux pump, PLoS One 9, e97903.

[6] Hagman, K. E., Lucas, C. E., Balthazar, J. T., Snyder, L., Nilles, M., Judd, R. C., and Shafer, W. M. (1997) The MtrD protein of Neisseria gonorrhoeae is a member of the resistance/nodulation/division protein family constituting part of an efflux system, Microbiology 143 (Pt 7), 2117–2125.

[7] Unemo, M., and Shafer, W. M. (2014) Antimicrobial resistance in Neisseria gonorrhoeae in the 21st century: past, evolution, and future, Clin Microbiol Rev 27, 587–613.

[8] Chitsaz, M., Booth, L., Blyth, M. T., O’Mara, M. L., and Brown, M. H. (2019) Multidrug Resistance in Neisseria gonorrhoeae: Identification of Functionally Important Residues in the MtrD Efflux Protein, mBio 10.

[9] Warner, D. M., Shafer, W. M., and Jerse, A. E. (2008) Clinically relevant mutations that cause derepression of the Neisseria gonorrhoeae MtrC-MtrD-MtrE Efflux pump system confer different levels of antimicrobial resistance and in vivo fitness, Mol Microbiol 70, 462–478.

[10] Lyu, M., Moseng, M. A., Reimche, J. L., Holley, C. L., Dhulipala, V., Su, C. C., Shafer, W. M., and Yu, E. W. (2020) Cryo-EM Structures of a Gonococcal Multidrug Efflux Pump Illuminate a Mechanism of Drug Recognition and Resistance, mBio 11.

[11] Eicher, T., Seeger, M. A., Anselmi, C., Zhou, W., Brandstatter, L., Verrey, F., Diederichs, K., Faraldo-Gomez, J. D., and Pos, K. M. (2014) Coupling of remote alternating-access transport mechanisms for protons and substrates in the multidrug efflux pump AcrB, Elife 3.

[12] Matsunaga, Y., Yamane, T., Terada, T., Moritsugu, K., Fujisaki, H., Murakami, S., Ikeguchi, M., and Kidera, A. (2018) Energetics and conformational pathways of functional rotation in the multidrug transporter AcrB, Elife 7.

[13] Murakami, S., Nakashima, R., Yamashita, E., Matsumoto, T., and Yamaguchi, A. (2006) Crystal structures of a multidrug transporter reveal a functionally rotating mechanism, Nature 443, 173–179.

[14] Du, D., Voss, J., Wang, Z., Chiu, W., and Luisi, B. F. (2015) The pseudo-atomic structure of an RND-type tripartite multidrug efflux pump, Biol Chem 396, 1073–1082.

[15] Eicher, T., Cha, H. J., Seeger, M. A., Brandstatter, L., El-Delik, J., Bohnert, J. A., Kern, W. V., Verrey, F., Grutter, M. G., Diederichs, K., and Pos, K. M. (2012) Transport of drugs by the multidrug transporter AcrB involves an access and a deep binding pocket that are separated by a switch-loop, Proc Natl Acad Sci U S A 109, 5687–5692.

[16] Su, C. C., Yin, L., Kumar, N., Dai, L., Radhakrishnan, A., Bolla, J. R., Lei, H. T., Chou, T. H., Delmar, J. A., Rajashankar, K. R., Zhang, Q., Shin, Y. K., and Yu, E. W. (2017) Structures and transport dynamics of a Campylobacter jejuni multidrug efflux pump, Nat Commun 8, 171.

[17] Nakashima, R., Sakurai, K., Yamasaki, S., Nishino, K., and Yamaguchi, A. (2011) Structures of the multidrug exporter AcrB reveal a proximal multisite drug-binding pocket, Nature 480, 565–569.

[18] Rouquette-Loughlin, C. E., Reimche, J. L., Balthazar, J. T., Dhulipala, V., Gernert, K. M., Kersh, E. N., Pham, C. D., Pettus, K., Abrams, A. J., Trees, D. L., St Cyr, S., and Shafer, W. M. (2018) Mechanistic Basis for Decreased Antimicrobial Susceptibility in a Clinical Isolate of Neisseria gonorrhoeae Possessing a Mosaic-Like mtr Efflux Pump Locus, mBio 9.

[19] Wang, Z., Fan, G., Hryc, C. F., Blaza, J. N., Serysheva, II, Schmid, M. F., Chiu, W., Luisi, B. F., and Du, D. (2017) An allosteric transport mechanism for the AcrAB-TolC multidrug efflux pump, Elife 6.

[20] McCormick, J. W., Ammerman, L., Chen, G., Vogel, P. D., and Wise, J. G. (2021) Transport of Alzheimer’s associated amyloid-beta catalyzed by P-glycoprotein, PLoS One 16, e0250371.

[21] Hanamoto, S., and Ogawa, F. (2019) Predicting the sorption of azithromycin and levofloxacin to sediments from mineral and organic components, Environ Pollut 255, 113180.

[22] Smolarchuk, C., Wensley, A., Padfield, S., Fifer, H., Lee, A., and Hughes, G. (2018) Persistence of an outbreak of gonorrhoea with high-level resistance to azithromycin in England, November 2014May 2018, Euro Surveill 23.

[23] Trott, O., and Olson, A. J. (2010) AutoDock Vina: improving the speed and accuracy of docking with a new scoring function, efficient optimization, and multithreading, J Comput Chem 31, 455–461.

[24] Webb, B., and Sali, A. (2016) Comparative Protein Structure Modeling Using MODELLER, Curr Protoc Protein Sci 86, 2 9 1–2 9 37.

[25] Kong, F. Y., Rupasinghe, T. W., Simpson, J. A., Vodstrcil, L. A., Fairley, C. K., McConville, M. J., and Hocking, J. S. (2017) Pharmacokinetics of a single 1g dose of azithromycin in rectal tissue in men, PLoS One 12, e0174372.

[26] ACE and JChem acidity and basicity Calculator, ACE UKY-4.0 ed., Marvin JS, ChemAxon, https://epoch.uky.edu/ace/public/pKa.jsp.

[27] Rostkowski, M., Olsson, M. H., Sondergaard, C. R., and Jensen, J. H. (2011) Graphical analysis of pH-dependent properties of proteins predicted using PROPKA, BMC Struct Biol 11, 6.

[28] McFarland, J. W., Berger, C. M., Froshauer, S. A., Hayashi, S. F., Hecker, S. J., Jaynes, B. H., Jefson, M. R., Kamicker, B. J., Lipinski, C. A., Lundy, K. M., Reese, C. P., and Vu, C. B. (1997) Quantitative structure-activity relationships among macrolide antibacterial agents: in vitro and in vivo potency against Pasteurella multocida, J Med Chem 40, 1340–1346.

[29] Jerse, A. E., Sharma, N. D., Simms, A. N., Crow, E. T., Snyder, L. A., and Shafer, W. M. (2003) A gonococcal efflux pump system enhances bacterial survival in a female mouse model of genital tract infection, Infect Immun 71, 5576–5582.

[30] Ng, L. K., and Martin, I. E. (2005) The laboratory diagnosis of Neisseria gonorrhoeae, Can J Infect Dis Med Microbiol 16, 15–25.

[31] Vargiu, A. V., Ramaswamy, V. K., Malvacio, I., Malloci, G., Kleinekathofer, U., and Ruggerone, P. (2018) Water-mediated interactions enable smooth substrate transport in a bacterial efflux pump, Biochim Biophys Acta Gen Subj 1862, 836–845.

[32] Alberts B, J. A., Lewis J, et al. (2002) Molecular Biology of the Cell. 4th Edition., (Figure 14–32, t. I. o. H.-D. T. i. B., Ed.), Garland, New York.

[33] Gaillard, P., Carrupt, P. A., Testa, B., and Boudon, A. (1994) Molecular lipophilicity potential, a tool in 3D QSAR: method and applications, J Comput Aided Mol Des 8, 83–96.

[34] Ramaswamy, V. K., Vargiu, A. V., Malloci, G., Dreier, J., and Ruggerone, P. (2018) Molecular Determinants of the Promiscuity of MexB and MexY Multidrug Transporters of Pseudomonas aeruginosa, Front Microbiol 9, 1144.

[35] Goddard, T. D., Huang, C. C., Meng, E. C., Pettersen, E. F., Couch, G. S., Morris, J. H., and Ferrin, T. E. (2018) UCSF ChimeraX: Meeting modern challenges in visualization and analysis, Protein Sci 27, 14–25.

[36] Dolinsky, T. J., Czodrowski, P., Li, H., Nielsen, J. E., Jensen, J. H., Klebe, G., and Baker, N. A. (2007) PDB2PQR: expanding and upgrading automated preparation of biomolecular structures for molecular simulations, Nucleic Acids Res 35, W522–525.

[37] Schulz, R., Vargiu, A. V., Collu, F., Kleinekathofer, U., and Ruggerone, P. (2010) Functional rotation of the transporter AcrB: insights into drug extrusion from simulations, PLoS Comput Biol 6, e1000806.

[38] Wadsworth, C. B., Arnold, B. J., Sater, M. R. A., and Grad, Y. H. (2018) Azithromycin Resistance through Interspecific Acquisition of an Epistasis-Dependent Efflux Pump Component and Transcriptional Regulator in Neisseria gonorrhoeae, mBio 9.

[39] Vargiu, A. V., and Nikaido, H. (2012) Multidrug binding properties of the AcrB efflux pump characterized by molecular dynamics simulations, Proc Natl Acad Sci U S A 109, 20637–20642.

[40] Humphrey, W., Dalke, A., and Schulten, K. (1996) VMD: visual molecular dynamics, J Mol Graph 14, 33–38, 27-38.

[41] Lee, J., Cheng, X., Swails, J. M., Yeom, M. S., Eastman, P. K., Lemkul, J. A., Wei, S., Buckner, J., Jeong, J. C., Qi, Y., Jo, S., Pande, V. S., Case, D. A., Brooks, C. L., 3rd, MacKerell, A. D., Jr., Klauda, J. B., and Im, W. (2016) CHARMM-GUI Input Generator for NAMD, GROMACS, AMBER, OpenMM, and CHARMM/OpenMM Simulations Using the CHARMM36 Additive Force Field, J Chem Theory Comput 12, 405–413.

[42] Jo, S., Kim, T., Iyer, V. G., and Im, W. (2008) CHARMM-GUI: a web-based graphical user interface for CHARMM, J Comput Chem 29, 1859–1865.

[43] Brooks, B. R., Brooks, C. L., 3rd, Mackerell, A. D., Jr., Nilsson, L., Petrella, R. J., Roux, B., Won, Y., Archontis, G., Bartels, C., Boresch, S., Caflisch, A., Caves, L., Cui, Q., Dinner, A. R., Feig, M., Fischer, S., Gao, J., Hodoscek, M., Im, W., Kuczera, K., Lazaridis, T., Ma, J., Ovchinnikov, V., Paci, E., Pastor, R. W., Post, C. B., Pu, J. Z., Schaefer, M., Tidor, B., Venable, R. M., Woodcock, H. L., Wu, X., Yang, W., York, D. M., and Karplus, M. (2009) CHARMM: the biomolecular simulation program, J Comput Chem 30, 1545–1614.

[44] Lomize, M. A., Pogozheva, I. D., Joo, H., Mosberg, H. I., and Lomize, A. L. (2012) OPM database and PPM web server: resources for positioning of proteins in membranes, Nucleic Acids Res 40, D370–376.

[45] Case, D. A., Cheatham, T. E., 3rd, Darden, T., Gohlke, H., Luo, R., Merz, K. M., Jr., Onufriev, A., Simmerling, C., Wang, B., and Woods, R. J. (2005) The Amber biomolecular simulation programs, J Comput Chem 26, 1668–1688.

[46] D.A. Case, I. Y. B.-S., S.R. Brozell, D.S. Cerutti, T.E. Cheatham, III, V.W.D. Cruzeiro, T.A. Darden, R.E. Duke, D. Ghoreishi, G. Giambasu, T. Giese, M.K. Gilson, H. Gohlke, A.W. Goetz, D. Greene, R Harris, N. Homeyer, Y. Huang, S. Izadi, A. Kovalenko, R. Krasny, T. Kurtzman, T.S. Lee, S. LeGrand, P. Li, C. Lin, J. Liu, T. Luchko, R. Luo, V. Man, D.J. Mermelstein, K.M. Merz, Y. Miao, G. Monard, C. Nguyen, H. Nguyen, A. Onufriev, F. Pan, R. Qi, D.R. Roe, A. Roitberg, C. Sagui, S. Schott-Verdugo, J. Shen, C.L. Simmerling, J. Smith, J. Swails, R.C. Walker, J. Wang, H. Wei, L. Wilson, R.M. Wolf, X. Wu, L. Xiao, Y. Xiong, D.M. York and P.A. Kollman (2019) AMBER 2019, University of California, San Francisco.

[47] Salomon-Ferrer, R., Gotz, A. W., Poole, D., Le Grand, S., and Walker, R. C. (2013) Routine Microsecond Molecular Dynamics Simulations with AMBER on GPUs. 2. Explicit Solvent Particle Mesh Ewald, J Chem Theory Comput 9, 3878–3888.

[48] Morris, G. M., Huey, R., Lindstrom, W., Sanner, M. F., Belew, R. K., Goodsell, D. S., and Olson, A. J.(2009) AutoDock4 and AutoDockTools4: Automated docking with selective receptor flexibility, J Comput Chem 30, 2785–2791.

[49] Grant, B. J., Rodrigues, A. P., ElSawy, K. M., McCammon, J. A., and Caves, L. S. (2006) Bio3d: an R package for the comparative analysis of protein structures, Bioinformatics 22, 2695–2696.

[50] O’Boyle, N. M., Banck, M., James, C. A., Morley, C., Vandermeersch, T., and Hutchison, G. R. (2011) Open Babel: An open chemical toolbox, J Cheminform 3, 33.

[51] Phillips, J. C., Braun, R., Wang, W., Gumbart, J., Tajkhorshid, E., Villa, E., Chipot, C., Skeel, R. D., Kale, L., and Schulten, K. (2005) Scalable molecular dynamics with NAMD, J Comput Chem 26, 1781–1802.

[52] McCormick, J. W., Vogel, P. D., and Wise, J. G. (2015) Multiple Drug Transport Pathways through Human P-Glycoprotein, Biochemistry 54, 4374–4390.

[53] Wise, J. G. (2012) Catalytic transitions in the human MDR1 P-glycoprotein drug binding sites, Biochemistry 51, 5125–5141.

[54] Wang, J., Wang, W., Kollman, P. A., and Case, D. A. (2006) Automatic atom type and bond type perception in molecular mechanical calculations, J Mol Graph Model 25, 247–260.

[55] Wang, J., Wolf, R. M., Caldwell, J. W., Kollman, P. A., and Case, D. A. (2004) Development and testing of a general amber force field, J Comput Chem 25, 1157–1174.

[56] D.A. Case, R. M. B., D.S. Cerutti, T.E. Cheatham, III, T.A. Darden, R.E. Duke, T.J. Giese, H. Gohlke, A.W. Goetz, N. H., S. Izadi, P. Janowski, J. Kaus, A. Kovalenko, T.S. Lee, S. LeGrand, P. Li, C., Lin, T. L., R. Luo, B. Madej, D. Mermelstein, K.M. Merz, G. Monard, H. Nguyen, H.T. Nguyen, I., Omelyan, A. O., D.R. Roe, A. Roitberg, C. Sagui, C.L. Simmerling, W.M. Botello-Smith, J. Swails, and R.C. Walker, J. W., R.M. Wolf, X. Wu, L. Xiao and P.A. Kollman. (2016) AMBER 2016, University of California, San Francisco.

[57] Baker, N. A., Sept, D., Joseph, S., Holst, M. J., and McCammon, J. A. (2001) Electrostatics of nanosystems: application to microtubules and the ribosome, Proc Natl Acad Sci U S A 98, 10037–10041.

[58] Dolinsky, T. J., Nielsen, J. E., McCammon, J. A., and Baker, N. A. (2004) PDB2PQR: an automated pipeline for the setup of Poisson-Boltzmann electrostatics calculations, Nucleic Acids Res 32, W665–667.

